# An antibody drug conjugate targeting a GSTA glycosite-signature epitope of mucin1 expressed by non-small cell lung cancer

**DOI:** 10.1101/2019.12.22.885566

**Authors:** Deng Pan, Yubo Tang, Jiao Tong, Chengmei Xie, Jiaxi Chen, Chunchao Feng, Patrick Hwu, Wei Huang, Dapeng Zhou

## Abstract

**Background:** Antibodies targeting abnormally glycosylated proteins have been ineffective in treating cancer. Antibody-drug conjugates are emerging as an efficient option, which allow specific delivery of drugs into tumors. We and others have dissected the abnormally glycosylated tandem repeat region of MUC1 glycoprotein as three site-specific glycosylated neoantigen peptide motifs (PDTR, GSTA, GVTS) for monoclonal antibody binding.

**Methods:** Internalization of monoclonal antibodies was studied by immunofluorescence staining and colocalization with lysosomal markers in live cells. Antibody positivity in tumor and peritumoral tissue samples were studied by immunohistochemistry. The efficacy of anti-MUC1 ADCs were evaluated with various cancer cell lines and mouse tumor xenograft model.

**Results:** We describe an anti-MUC1 ADC by conjugating GSTA neoantigen-specific 16A with monomethyl auristatin E (MMAE). 16A-MMAE showed potent antitumoral efficacy with IC_50_ ranging from 0.2 to 49.4 nM toward multiple types of cancer cells. *In vivo*, 16A-MMAE showed dose-dependent inhibition of tumor growth in mouse xenograft of NCI-H838 NSCLC cell line, with minimum effective dose at 1 mg/kg. At the dose of 3 mg/kg, 16A-MMAE did not cause significant toxicity in a transgenic mouse expressing human MUC1.

**Conclusions:** The high antitumoral efficacy of 16A-MMAE suggest that aberrant glycosylated MUC1 neoantigen is a target with high positivity in multiple cancer types for ADC development. Personalized therapy may be achieved by development of glycosite-specific antibody-drug conjugates.

## Introduction

Mucin-1 (MUC1), also known as EMA, PEM or CA15-3 antigen, is a transmembrane glycoprotein that has been studied as a significant target for tumor immunotherapy.^1, 2^ In healthy cells, MUC1 is a heavily O-glycosylated protein. The extracellular portion contains a variable number of tandem repeats (VNTR), and the number of tandem repeats range from 20 to 120. Each TR consists of 20 amino acids and there are five potential O-glycosylation sites.^3, 4^ MUC1 is expressed in almost all epithelial cancers.^5–8^ Tumor MUC1 differs from normal MUC1 by abnormal, truncated glycosylation. The truncated glycosylation forms glycopeptide epitopes that can be recognized by specific antibodies. Since such glycopeptide epitopes are tumor specific, they may represent potential targets for therapeutic antibodies. However, monoclonal antibody therapeutics targeting MUC1 has not shown efficacy in clinical trials.^9, 10^ It was hypothesized MUC1 subunit containing the tandem repeats circulates at high levels in cancer patients and acts as a “sink” precluding delivery of antibodies to the tumor cell surface. However, significant inhibition of circulating MUC1 on antibody-dependent cellular cytotoxicity was only observed in patient serum with MUC1 levels above 100 U/ml.^11^ High levels of circulating autoantibodies against both the cancer-specific isoform of MUC1 and the non-glycosylated signal peptide domain of MUC1 (up to 200 μg/ml) were reported in human cancers.^12^ Several groups reported that autoantibodies toward a single glycopeptides epitope in cancer patients is highly variable.^13^ Some researchers have reported that autoantibodies to aberrantly glycosylated MUC1 in early stage breast cancer are associated with a better prognosis.^14^ Whether autoantibodies toward MUC1 affect the efficacy and specific targeting of antibody drug remain unclear. There is no data on affinity of autoantibodies toward MUC1. Clearly, for development of therapeutic antibodies, high-affinity is a critical criteria.

Recent efforts have been focused on evaluating MUC1’s potential as candidates for antibody-drug conjugates. An antibody-drug conjugate (ADC) consists of three components: an antibody, an antitumoral agent and a linker. The first ADC approved by FDA was gemtuzumabozogamicin (Mylotarg), a humanized anti-CD33 IgG4 antibody conjugated to calicheamicin, a potent cytotoxic agent that causes double-strand DNA breaks.^15^ It was used to treat patients with relapsed acute myeloid leukemia. There are currently two other FDA approved ADCs in clinic. Brentuximabvedotin (Adcetris) is an anti-CD30 antibody linked to a MMAE, an antimitotic agent which inhibits cell division by blocking the polymerisation of tubulin. It was approved for the treatment of relapsed or refractory systemic anaplastic large cell lymphoma or Hodgkin’s lymphoma.^16^ Trastuzumabemtansine (Kadcyla) is an anti-Her2 antibody linked to the tubulin inhibitor maytansine derivative DM1 (T-DM1). It is used in advanced Her2-positive breast cancer patients.^17^ Furthermore, more than 30 ADCs are in clinical development, targeting a wide range of blood tumors and solid carcinomas.^18^

We and others have dissected the abnormally glycosylated TR region as neoantigen peptide motifs (PDTR, GSTA, GVTS) for monoclonal antibody binding.^2, 19^ In this study, we screened monoclonal antibodies specific to above three neoantigen peptide motifs and synthesized antibody-MMAE for treatment of cancer.

## Materials and Methods

### Cell lines and reagents

Human tumor cell lines NCI-H838 (H838), NCI-H2030 (H2030), NCI-H1650 (H1650), NCI-H1975 (H1975), NCI-H23 (H23), NCI-H520 (H520), NCI-H460 (H460), NCI-H292 (H292), NCI-H1229 (H1229), A549, PC9, MCF-7, SKBR3, PANC-1, CFPAC1, N87, HGC-27, H8910, SKOV3, ES2, Hey and KGN (obtained from the America Type Culture Collection, ATCC) were cultured at 37 °C with 5% CO_2_ in RPMI-1640 or DMEM media (Life Technologies) supplemented with 10% fetal bovine serum (FBS, Life Technologies). The Endo-S was expressed in *E. coli* following the reported procedure.^20^ MMAE and Fmoc-ValCit (valine-citrulline)-PAB-PNP were purchased from Levena Biopharma (Nanjing, China). Other chemical reagents and solvents were purchased from Sinopharm Chemical Reagent Co. (Shanghai, China) or Sigma-Aldrich and used without further purification. The MAbPac RP column (4 μm, 3.0 × 100 mm) was purchased from ThermoFisher. Nuclear magnetic resonance (NMR) spectra were measured on a Varian-MERCURY Plus-500 instrument. ESI-HRMS spectra were measured on an Agilent 6230 LC-TOF MS spectrometer.

### Confocal microscopy

H838 cells were seeded on glass covers lips in 3.5 cm dishes (1.5 ×10^5^ cells per well) and cultured at 37 °C with 5% CO_2_ in RPMI-1640 with 10% fetal bovine serum for 24 hours. Cells were incubated with 2 μg/ml cy5-labelled 16A mAb for 3.5 hours, and subsequently incubated with 75 nM lysosome fluorescent probe (LysoTracker Red DND 99, ThermoFisher) for 30 min. Cells were washed, fixed, and observed with a confocal laser scanning microscope (Nikon A1R, Japan).

### Internalization of 16A antibody

To measure the internalization of antibody, multiple sets of 2 ×10^5^ H838 cells were first incubated with unlabeled 16A mAb at a saturating concentration of 5 μg/ml. The negative control group without internalization was kept on ice for 150 min. For internalization experiments, other sets of antibody-coated cells were incubated at 37 °C to allow antibody internalization for different time periods (30, 60, 90, 120 and 150 min). At the end of incubation, cells were washed with ice cold PBS buffer and then stained with a PE-labelled anti-mouse IgG secondary antibody (Southern Biotech, Birmingham, AL) at 1 μg/ml concentration for 30 min on ice. After 3 washes with PBS, the cells were harvested and analyzed by flow cytometry to calculate mean fluorescent intensity (MFI). The amount of 16A antibody internalized into cells at each time point was determined by the percentage of MFI decrease as compared to control cells that were incubated at 4 °C for 150 min.

### Flow cytometry staining of cancer cell lines

Cell surface expression of MUC1 was assessed by flow cytometry staining. The cells were washed with 2% BSA in PBS and then incubated with the 16A^21^, SM3 or C595 antibody (Abcam, US) at 5 μg/ml for 30 min at 4 °C. After washing, cells were incubated with PE-labelled anti-mouse IgG (1 μg/ml) for 30 min at 4 °C. After washing, cells were analyzed by FACS Calibur (BD company) and the data were analyzed using FlowJo software (version 7.6).

### Immunohistochemistry

Tumor tissue array slides were from Crownbio, China. Immunohistochemistry was performed by Bond RX automatic IHC&ISH machine, Leica. In brief, paraffin-embedded tissue samples were treated by Dewax solution and antigen retrieval buffer sequentially, followed by staining with primary and secondary antibodies. All of the staining was scanned with NanoZoomer Image system. The intensity of IHC staining were scored at four levels, 0 (negative), 1 (weak staining), 2 (medium staining), 3 (strong staining). The percentages of tumor cells at different intensity levels are evaluated.

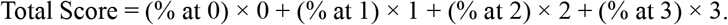

### Synthesis of NHS-ValCit-PAB-MMAE

Antibody-MMAE was synthesized as previously described.^22^ NH_2_-ValCit-PAB-MMAE (29.7 mM in 300 μl DMF) was added gradually (30 μl every 15 min) to a solution of disuccinimidyl glutarate (DSG, 53.4 mM) prepared in a mixture (1:1 volume) of DMF/phosphate buffer (50 mM, pH 7.5). The reaction mixture was stirred at room temperature for 3 hours and monitored by RP-HPLC. The product was purified by preparative HPLC to give a white powder (9.1 mg, 76.5%). 1H NMR (500 MHz, DMSO-d6) δ 8.14 (d, J = 7.5 Hz, 1H), 7.89 (d, J = 8.6 Hz, 2H), 7.59 (d, J = 8.1 Hz, 2H), 7.32 (d, J = 7.8 Hz, 2H), 7.28 (dd, J = 7.5, 2.7 Hz, 2H), 7.17 (s, 1H), 5.98 (s, 1H), 5.04 (dd, J = 31.4, 17.1 Hz, 3H), 4.50 (d, J = 5.9 Hz, 1H), 4.44 (d, J = 6.6 Hz, 1H), 4.39 (d, J = 8.3 Hz, 1H), 4.31-4.26 (m, 1H), 4.26-4.18 (m, 2H), 4.06-3.93 (m, 3H), 3.61-3.55 (m, 2H), 3.25 (d, J = 7.7 Hz, 3H), 3.20 (d, J = 12.5 Hz, 3H), 3.12 (s, 1H), 2.98 (s, 1H), 2.87 (dd, J = 18.5, 5.7 Hz, 3H), 2.82 (s, 3H), 2.69 (d, J = 7.5 Hz, 2H), 2.42 (d, J = 16.2 Hz, 2H), 2.36-2.23 (m, 4H), 2.17-2.08 (m, 2H), 2.01-1.68 (m, 10H), 1.63-1.47 (m, 4H), 1.07-0.98 (m, 6H), 0.91-0.74 (m, 21H). 13C NMR (126 MHz, DMSO-d6) δ 171.42, 171.16, 170.54, 170.20, 168.71, 158.82, 143.61, 128.11, 127.74, 127.68, 126.68, 126.60, 126.40, 126.35, 85.37, 74.74, 60.87, 58.12, 57.55, 57.10, 54.93, 54.06, 53.10, 47.15, 46.19, 43.70, 43.15, 38.55, 33.58, 31.48, 30.37, 29.87, 29.63, 29.25, 26.78, 25.41, 25.30, 24.29, 23.07, 20.56, 19.19, 18.88, 18.12, 15.41, 15.24, 14.96, 10.26. HRMS Calcd for [M+H]+ 1334.7612, found 1334.7586. [M+Na]+ 1356.7931, found 1356.7399.

### Preparation of lysine-linked antibody-drug conjugates

16A monoclonal antibody (1 mg/ml) and NHS-ValCit-PAB-MMAE (1.5 mM) in phosphate buffer (pH 8.0, 50 mM) containing 4-5% DMSO was incubated at 37 °C for 2 hours. The reaction mixture was immediately subject to protein-A affinity chromatography column for purification. Before loading the ADC sample, the protein A-agarose column was pre-washed with a glycine-HCl (100 mM, pH 2.5, 5 column volume) and pre-equilibrated with phosphate buffer (50 mM, pH 8.0, 5 column volume). After loading the ADC reaction mixture, the column was washed with phosphate buffer (50 mM, pH 8.0, 5 column volume) and glycine-HCl (20 mM, pH 5.0, 3 column volume) successively. Then the bound ADC was eluted with glycine-HCl (100 mM, pH 2.5, 5 column volume) followed by neutralization to pH 7.5 with glycine-HCl (1 M, pH 8.8) immediately. The fractions containing the target ADC were combined and concentrated by centrifugal filtration through a 10 kDa cut-off membrane.

### High performance liquid chromatography (HPLC) of 16A-MMAE

Analytical RP-HPLC was performed on LC3000 (analytic) instrument (Beijing ChuangXinTongHeng Ltd, China) with a C18 column (5 μm, 4.6 ×150 mm) at 40 °C. The column was eluted with a linear gradient of 2-90% acetonitrile containing 0.1% TFA for 30 min at a flow rate of 1 ml/min. Preparative HPLC was performed on LC3000 instrument with a preparative column (Waters, C18, OBD, 5 μm, 19 ×250 mm). The column was eluted with a linear gradient of aqueous acetonitrile containing 0.1% TFA at a flow rate of 10 ml/min.

### Liquid chromatography mass spectrometry (LC-MS)

ESI-MS spectra of small molecules were measured on an Agilent 6230 LC-TOF MS spectrometer. The small molecules were analyzed using a short guard column and eluted with 70% methanol containing 0.1% formic acid. The mass spectra of small molecules were recorded in the mass range of 200-3000 or 600-2000 under a high resolution mass-spec mode (HRMS, standard 3200 m/z, 4 GHz). Key source parameters: a drying nitrogen gas flow of 11 L/min; a nebulizer pressure of 40 psi; a gas temperature of 350 °C; a fragmentor voltage of 175 V; a skimmer voltage of 65 V; and a capillary voltage of 4000 V.

LC-MS spectra of antibodies and ADCs were measured on the same MS spectrometer (Agilent 6230) with a THERMO MAbPac RP column (4 μm, 3.0 ×100 mm) at 80 °C. The column was eluted with an isocratic mobile phase of 20% acetonitrile (Buffer B) and 80% water containing 0.1% formic acid (Buffer A) for the first 3 min at a flow rate of 0.4 ml/min, then it was successively eluted at the same flow rate with a linear gradient of 20-50% acetonitrile for additional 2.5 min, an isocratic 50% acetonitrile for 2 min, another linear gradient of 50-90% acetonitrile 0.5 min, and an isocratic 90% acetonitrile for 2 min. The mass spectra of antibodies were collected under the extended mass range mode (high 20 000 m/z, 1 GHz) in the mass range of 800-5000. Key source parameters: a drying nitrogen gas flow of 11 L/min; a nebulizer pressure of 60 psi; a gas temperature of 350 °C; a fragmentor voltage of 400 V; a skimmer voltage of 65 V; and a capillary voltage of 5000 V. The multiple charged peaks of the antibody were deconvoluted using the Agilent MassHunter Bioconfirm software (deconvolution for protein, Agilent technology) with the deconvolution range from 20 kDa to 160 kDa; other parameters were set at default values for protein deconvolution. The TOF was calibrated over the range 0-6000 m/z using Agilent ESI calibration mix solution before analysis. The peak of MS 922 is the internal standard for calibration.

### Pharmacokinetic studies of ADC

All animal studies were approved by the animal care and use committee of Tongji University. All experiments were carried in SPF housing facilities. Groups of mice (n=3, C75BL/6 strain, 8-week-old) were injected intravenously with a single dose of non-conjugated 16A mAb or 16A-MMAE at 5 mg/kg (mean body weight of mice being 20 g). Whole blood sample was collected from tail vein at various time points (0.25, 6, 24, 48, 96, 144, and 192 hours post dose). Serum was separated by centrifugation and stored at −80 °C until analysis.

The antibody drug concentrations were analyzed by ELISA. 96-well plates were coated with 1.5 μg/ml streptavidin (S4762, Sigma) at 4 °C overnight. Plate was blocked with 1% bovine serum albumin in PBS at 37 °C for one hour, then incubated with 2 μg/ml biotinylated MUC1 glycopeptide antigen at 37 °C for 1 hour. After washing, the mouse plasma samples and calibration standards of 16A/16A-MMAE (serial concentration: 400 to 3.125 ng/ml) were added. The binding reaction of plasma antibody drug was for 1 hour at 37 °C. After washing, the plate was incubated with horseradish peroxidase (HRP)-conjugated anti-mouse IgG at 37 °C in 1 hour, and visualized by adding 100 μl of TMB substrate solution for 30 min at room temperature followed by 100 μl of terminating solution. The absorbance was measured at 450 nm. Antibody drug concentrations and pharmacokinetics parameters were calculated using the PK solver software.

### Cytotoxicity assay

The 3-(4,5-dimethyl-2-thiazolyl)-2,5-diphenyl-2-H-tetrazolium bromide (MTT) method was used to measure the *in vitro* efficacy of the ADCs. Briefly, cells were plated at 5000 cells per well in 96-well plates at 37 °C and 5% CO_2_ for overnight. The ADC samples were serially diluted from 100 to 10e-4 μg/ml in culture medium. Cell viability was assessed after 72 hours by the MTT method. IC_50_ was determined by GraphPad Prism version 5 (GraphPad Software, Inc. San Diego, CA).

### Efficacy of 16A-MMAE in mouse xenograft tumor models

Groups of Balb/c nu/nu mice (6-week-old, female) were inoculated with 1×10^7^ H838 cells in 200 μl of PBS on the right flank. Tumor size was measured starting on day 8 (day 7 after first injection) and then every 2-3 days. The longest length (a) and the length perpendicular to the longest length (b) were used in the formula V = ½ ×a ×(b)^2^ to obtain the tumor volume in mm^3^. Mice were randomized into different treatment groups (n =5) when tumors reached a size between 150 and 250 mm^3^, and treated by antibody drugs.

The different treatment groups include 16A (15 mg/kg), 16A-MMAE (0.5, 1, 3, 5 and 15 mg/kg), and vehicle PBS, respectively. Mice received one or two doses of ADCs (second dose was 48 hours after first dose), non-conjugated antibodies, or vehicle PBS administered by intravenous injection. Statistic analysis was performed with GraphPad Prism version 5 (GraphPad Software, Inc. San Diego, CA).

### 16A-MMAE toxicity in hMUC1 transgenic mice

Groups of hMUC1-Tg mice^22^ (024631-C57BL/6-Tg(MUC1)79.24Gend/J, 8-week-old, n=6 per group, 3 males and 3 females, from Jackson Laboratory) were treated by 16A-MMAE via tail vein injection at a single dose of 0, 3, 15, or 30 mg/kg. Systemic tissue toxicity was examined at the Shanghai Institute of Materia Medica’s Center for Drug Safety Evalution and Research. Clinical pathology parameters were assessed on days 3, 14, and 28. Tissue samples of heart, liver, spleen, lung, kidney, gastric, pancreatic, and small intestine from ADC treated hMUC1 transgenic mice were fixed in 4% formaldehyde and embedded in paraffin blocks. Tissue sections were stained with hematoxylin and eosin.

### Efficacy of 16A-MMAE on MUC1-expressing syngeneic tumor in hMUC1 transgenic mice

A B16-OVA-hMUC1 cell line was generated by stable transfection of B16-OVA cell line^23^ with a pcMV3-hygro(R) plasmid (Sino Biological, China) encoding human MUC1 gene. Groups of hMUC1-Tg mice^22^ (024631-C57BL/6-Tg(MUC1)79.24Gend/J, 8-week-old, from Jackson Laboratory) were inoculated with 1×10^5^ B16-OVA-hMUC1 cells in 200 μl of PBS on the right flank. Mice were randomized into different treatment groups (n = 4) when tumors reached a size about 100 mm^3^. The different treatment groups include 16A-MMAE (3 and 10 mg/kg), and vehicle PBS, respectively. Mice received two doses of 16A-MMAE (second dose was 48 hours after first dose), or vehicle PBS administered by intravenous injection. Statistic analysis was performed with GraphPad Prism version 5 (GraphPad Software, Inc. San Diego, CA).

## Results

### Internalization and delivery of 16A antibody to lysosomes

We have screened a panel of mAbs specific to abnormally glycosylated MUC1 peptide motifs (PDTR, GSTA, GVTS) and focused on 16A mAb.^21^ The cy5-labelled 16A was initially localized on the cell surface after 30 min incubation at 37 °C (Fig. 1A, green). After 4 hours incubation at 37 °C, 16A was effectively internalized through endocytosis. The intracellular 16A co-localized with LysoTrack Red DND 99, a red-fluorescent dye for labeling and tracking acidic organelles in live cells, indicating that 16A was internalized and transported to the lysosomes (Fig. 1A, yellow). Lung cancer cell line H838 internalized 16A antibody within 150 min (Fig. 1B).

**Fig 1.**
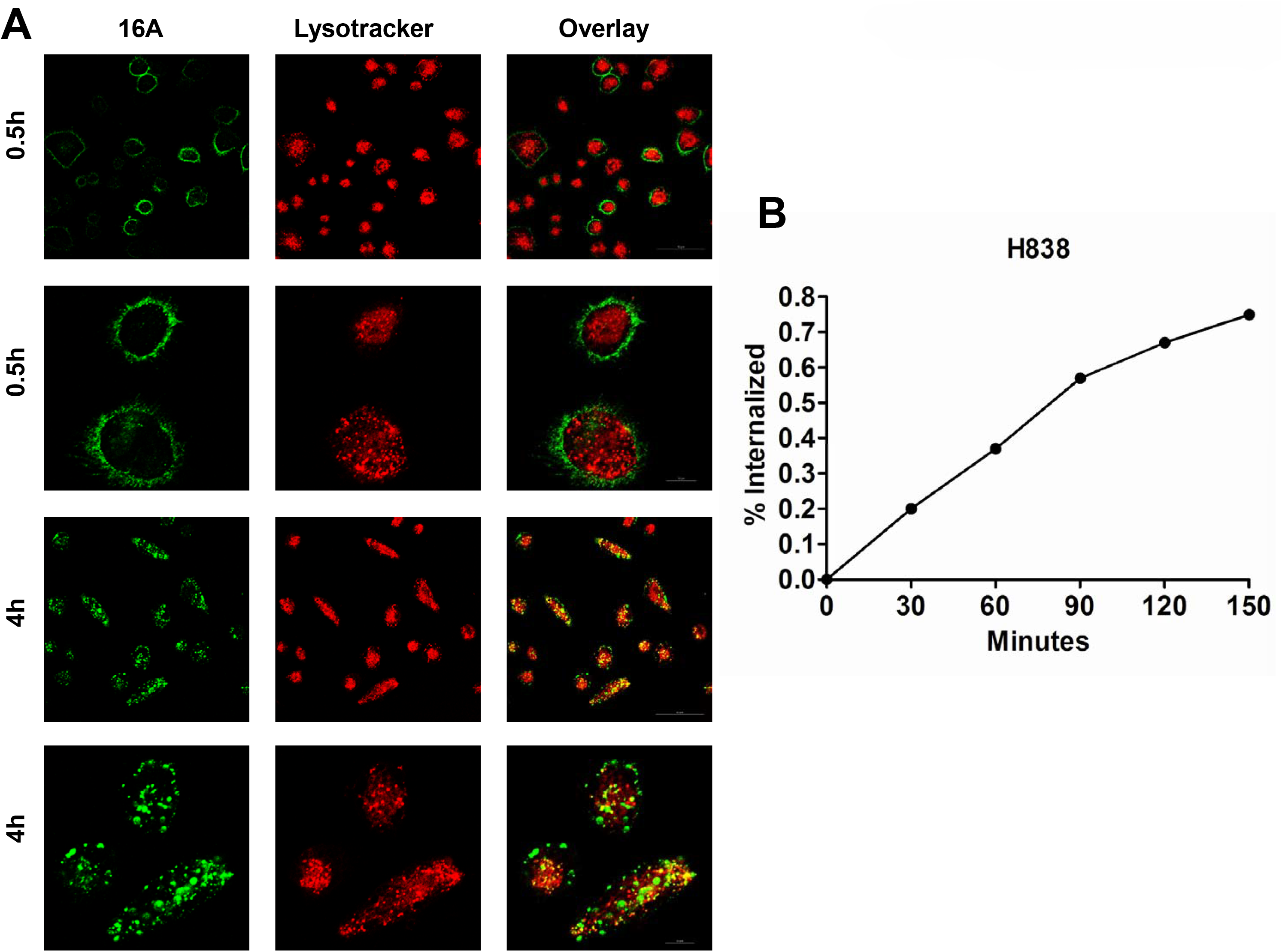
Internalization of 16A antibody. **(A)** Fluorescence-labeled 16A was incubated with H838 cells for 0.5 or 4 hours at 37 °C. After 0.5 hour, 16A staining was located to the cellular membrane. Four hours later, the antibody was endocytosed and co-localized with LysoTrack Red, a marker for acidic organelles in live cells. **(B)** Time course of 16A antibody internalization by lung cancer cell line H838 was measured by flow cytometry.

### Positivity of 16A antibody epitope in cancer cells and tissues

By flow cytometry analysis, we found that 16A antibody which specifically binds to GSTA neoantigen as we previously reported^21^ showed high positivity in 11 NSCLC cell lines (Fig. 2A). In contrast, SM3 and C595, two antibody clones which specifically bind to PDTR neoantigen epitope, showed extremely low positivity in NSCLC cell lines. We further confirmed this finding by immunohistochemistry staining in consecutive sections of tumor tissues from NSCLC patients. Tumors from same patients showed strong staining to 16 antibody, but are negative for SM3 or C595 (Fig. 2B, 2C and 2D).

**Fig 2.**
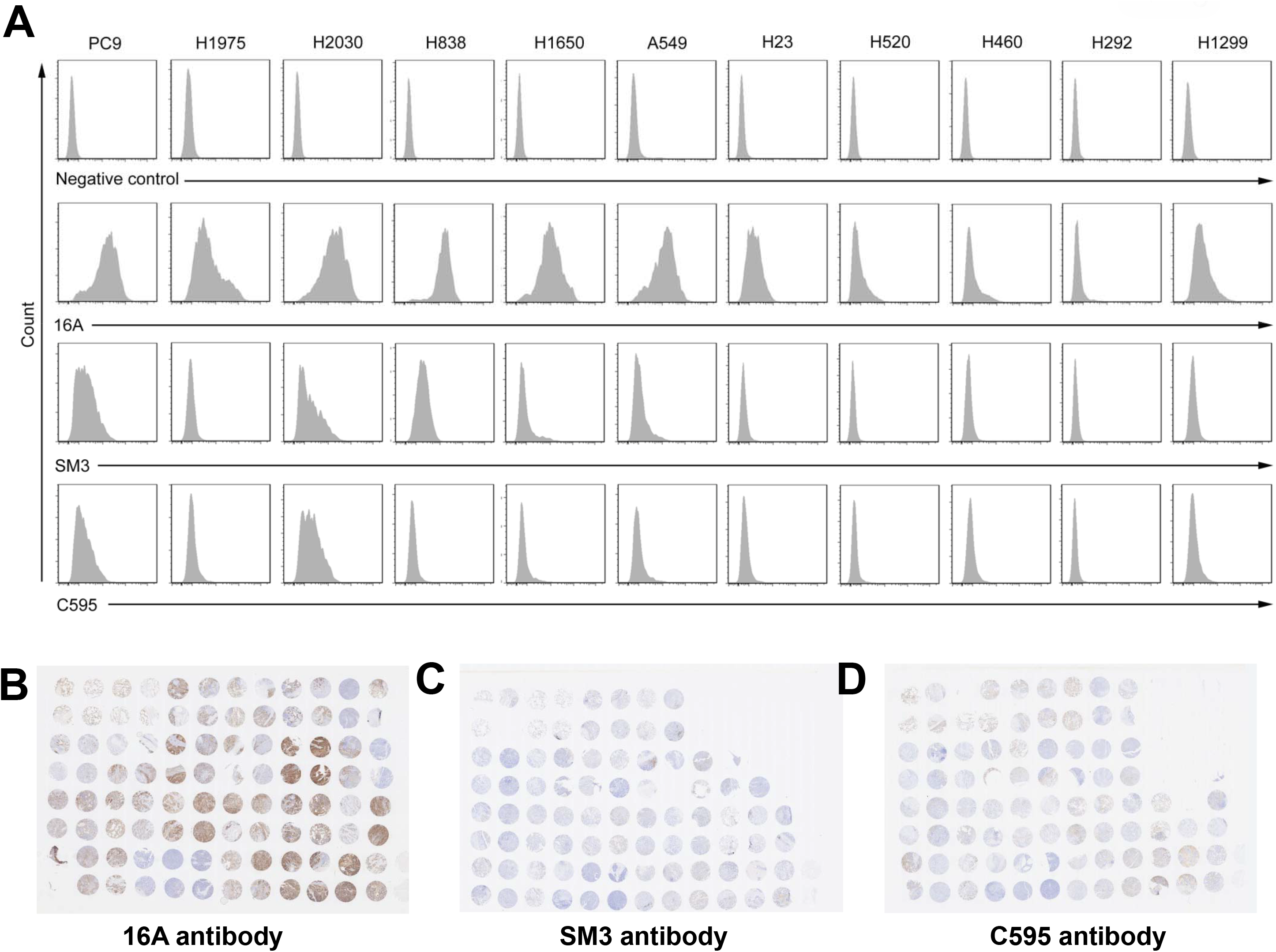
Staining of NSCLC cells and tissues by 16A (specific to GSTA neoantigen), SM3 and C595 (specific to PDTR neoantigen). **(A)** NSCLC cell lines were stained by 16A, SM3 and C595 antibodies. **(B)** Tissue array slides containing consecutive tissue sections from same NSCLC patients were stained by 16A, SM3 and C595 antibodies.

We and others previously reported findings that antibodies specific to tumor MUC1 preferentially bind to cell surface of tumor cells, but not heatlhy cells.^2^ IHC staining showed strong binding of 16A mAb to lung (Fig. 3A and Fig. S1), breast, TNBC (Fig. 3A and Fig. S2) and gastric (Fig. S3) cancer tissues. Positivity of 16A mAb staining was more than 60% in lung, breast, TNBC (Fig. 3B and Table 1) and gastric cancers (Fig. S3 and Table 1). Weak binding was found in colon and rectum cancer tissues by 16A antibody (Fig. S3 and Table 1). Significant higher expression was found in lung adenocarcinoma and squamous carcinoma as compared to peritumoral tissues (Fig. S4).

**Table 1.**
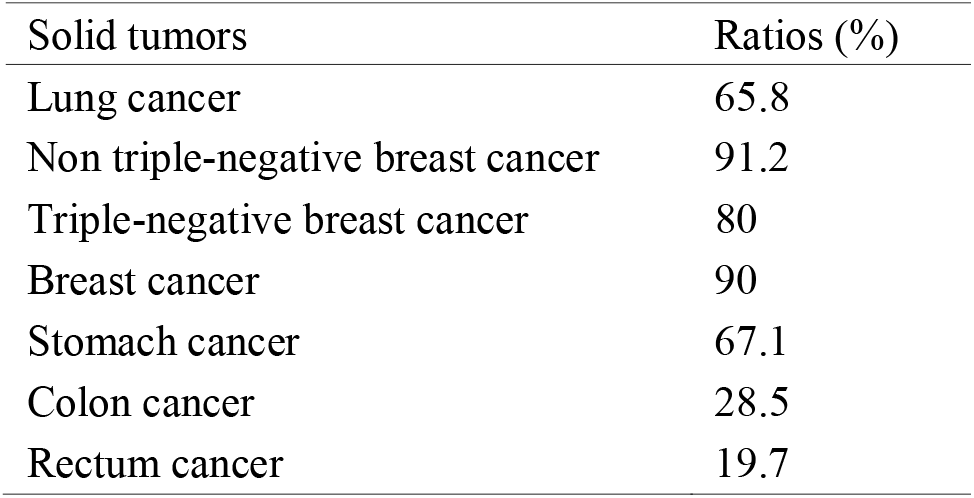
Positivity of 16A staining in solid tumors.

**Fig 3.**
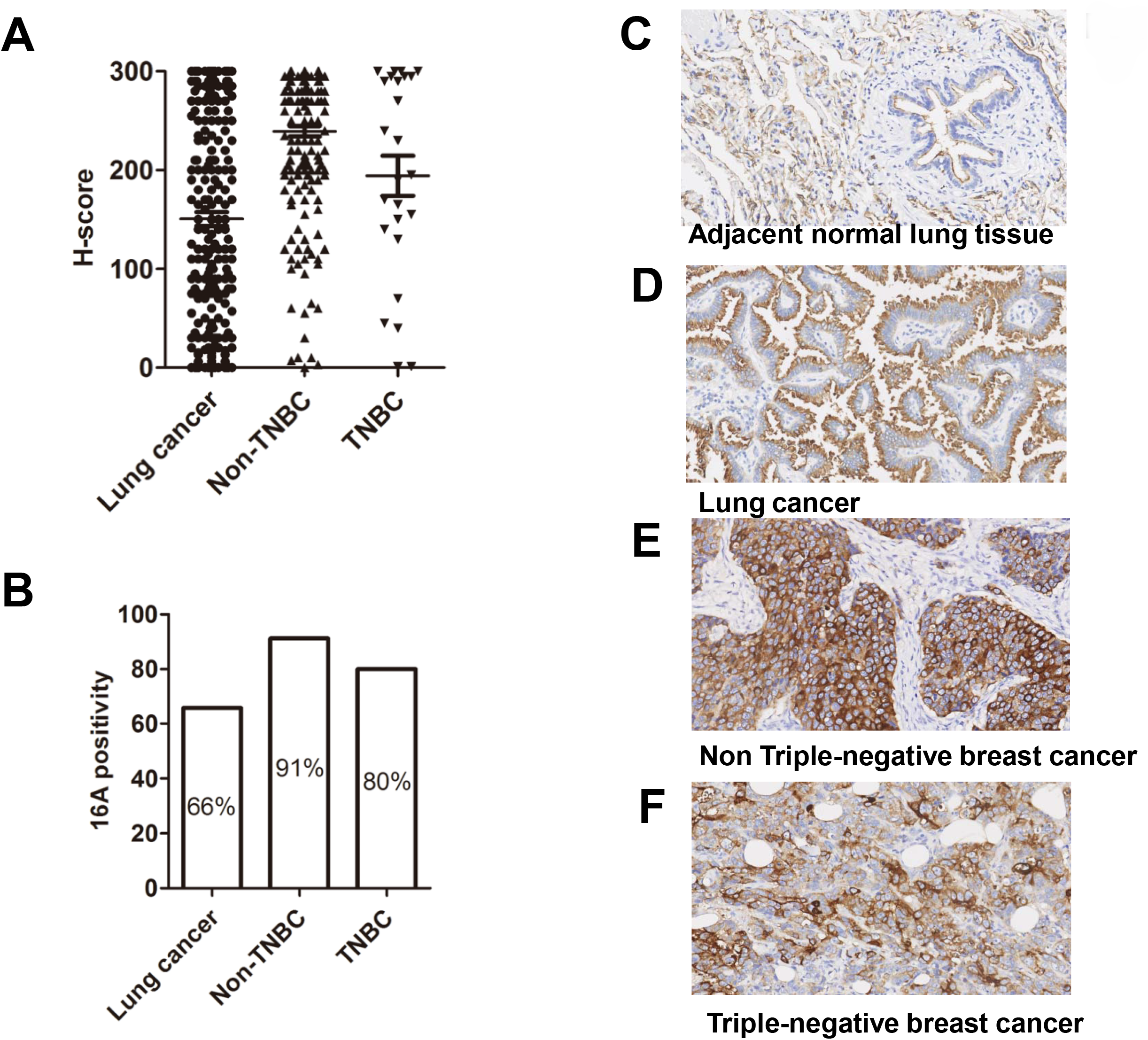
The expression of aberrant glycosylated MUC1 peptide motif in lung and breast cancer tissues as stained by 16A antibody. **(A)** H-score of cancer tissue array stained with 16A. The intensity of IHC staining were scored at four levels, 0 (negative), 1 (weak staining), 2 (medium staining), 3 (strong staining). The percentages of tumor cells at different intensity levels are evaluated. Total Score = (% at 0) ×0 + (% at 1) ×1 + (% at 2) ×2 + (% at 3) ×3. **(B)** 16A positivity in lung cancer, breast cancer and TNBC samples. **(C,D,E,F)** Representative photographs of MUC1 immunostaining inperitumoral (**C**), lung cancer (**D**), Non-TNBC (**E**) and TNBC (**F**) tissue by 16A antibody (original magnification ×200). Positive was defined as ≥30% of tumor with staining ≥2+.

16A staining was found in cytoplasm of peritumoral cells (Fig. 3C), but not cell surface. In contrast, strong staining by 16A was found on cell surface of tumor tissue cells (Fig. 3D). Moreover, strong staining by 16A was found in both cytoplasm and cell surface of breast and TNBC tissue cells (Fig. 3E and 3F).

Normal tissues including thymus, tonsil, spleen, cerebellum, pituitary, ovary, liver, paranephros, testis, intestine, cervix, salivary gland, bone marrow, cerebral cortex, bladder, striated muscle, and heart showed weak or no binding to 16A (Fig. S5). The staining to normal tissues are on cytoplasm of cells.

### Drug antibody ratio of 16A-MMAE

MMAE, a synthetic analog of the natural product dolastatin 10, originally isolated from sea hare, is a potent inhibitor of tubulin polymerization.^24^ We synthesized 16A-MMAE using a cleavable ValCit dipeptide linker connecting a payload with a *p*-aminobenzyloxycarbonyl (PABC) group (Fig. 4A). ValCit linkers are selectively cleaved by lysosomal enzymes upon internalization of 16A-MMAE by cancer cells, resulting in release of MMAE (Fig. 4B). LC-MS analysis was used to determine the average drug antibody ratio (DAR) of 16A-MMAE. The conjugation mixture contains 16A conjugates with 0, 1, 2, 3, 4, 5, and 6 drugs per antibody, and the average DAR of 16A-MMAE was 3.11 (Fig. 4C). The heterogeneity of DAR values is a common property of lysine-linked ADCs, including the T-DM1 reported in references.^25,26^ In our experiment, we employed consistent conjugation conditions (temperature, solvent, pH, concentration, and ratio of materials). Furthermore, we also combined the real-time DAR monitoring as we previously reported,^27^ to avoid the batch-to-batch inconsistency.

**Fig 4.**
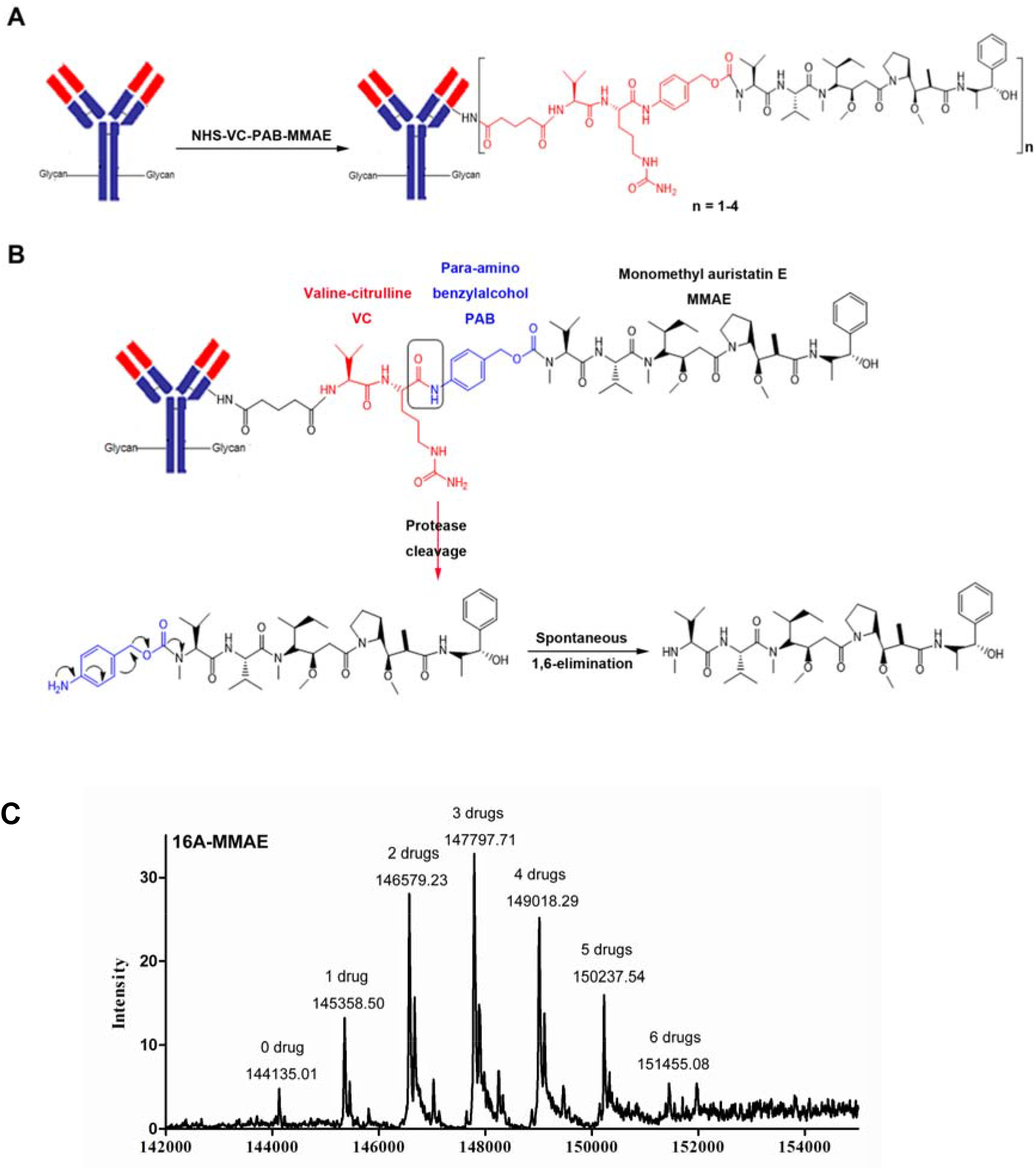
Preparation of ValCit-MMAE ADC and drug antibody ratio of 16A-MMAE. **(A)** The scheme for chemical synthesis of antibody-ValCit-MMAE. **(B)** The scheme of ValCit linker cleavage by the cathepsin B after the ADC is internalized by tumor cells. The activated MMAE drug is formed by spontaneous1,6-elimination. **(C)** LC-MS analysis of 16A-MMAE. The average drug antibody ratio is 3.11.

### Pharmacokinetics of 16A-MMAE

To determine the serum half-life of non-conjugated 16A antibody and 16A-MMAE in C57BL/6 mice, we intravenously injected 16A or 16A-MMAE and measured the serum antibody concentration at the different time points (0.25, 6, 24, 48, 96, 144, and 192 hours) (Fig. 5). The non-conjugated 16A antibody showed a serum half-life of 207.00 hours, and the 16A-MMAE showed a shorter serum half-life (144.22 hours, Table S1). The clearance and mean residence time are consistent with the serum half-life.

**Fig 5.**
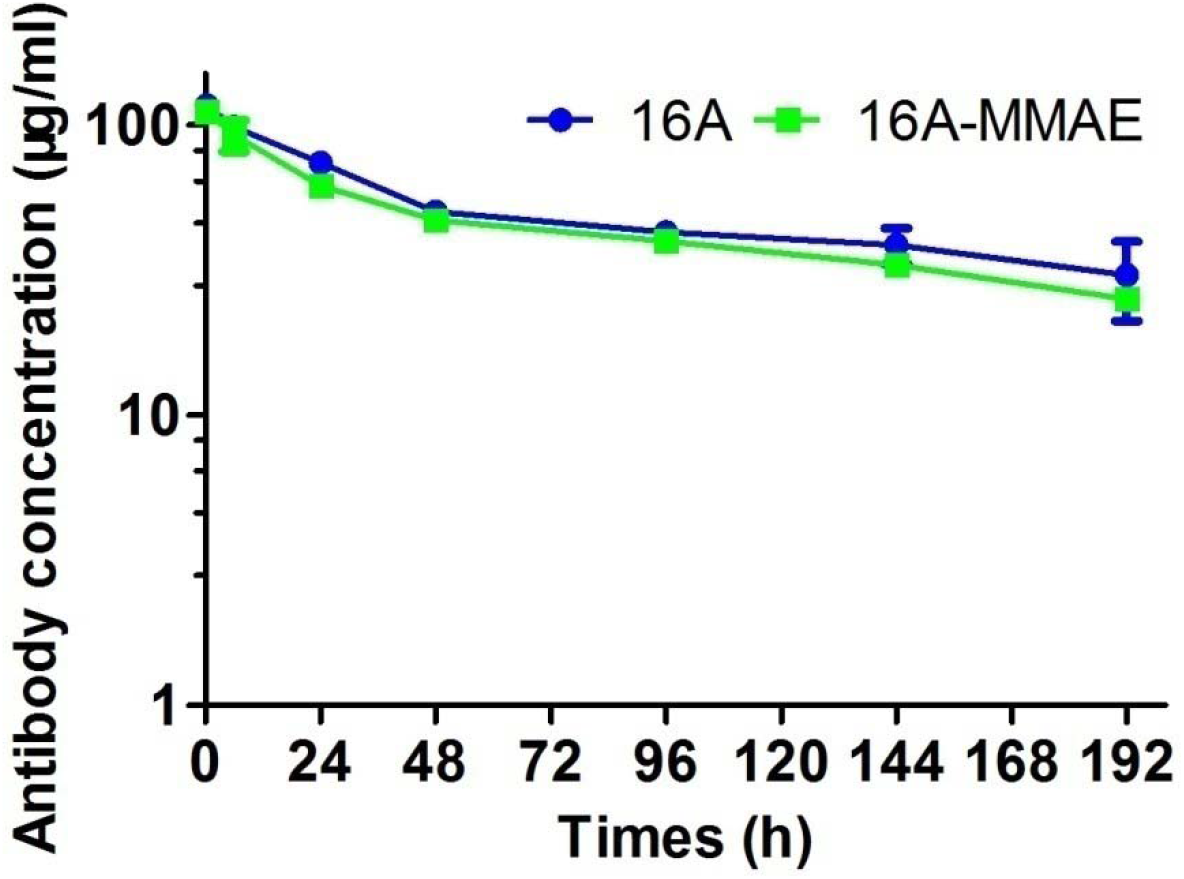
Pharmacokinetic profiles of non-conjugated 16A and 16A-MMAE *in vivo*. C57BL/6 mice were injected intravenously with a single dose of non-conjugated 16A mAb or 16A-MMAE ADC at 5 mg/kg. Serum antibody concentration was measured by ELISA at different time points.

### IC50 of 16A-MMAE

The cytotoxicity assay showed that 16A-MMAE exhibited strong tumor cell killing toward most of the lung, breast, pancreatic, gastric and ovarian (Fig. S6) cell lines. The IC_50_ of 16A-MMAE was shown in Table 2. Flow cytometry results showed that most of lung, breast, pancreatic and gastric (Fig. S6) cancer cells could be strongly stained by 16A antibody.

**Table 2.**
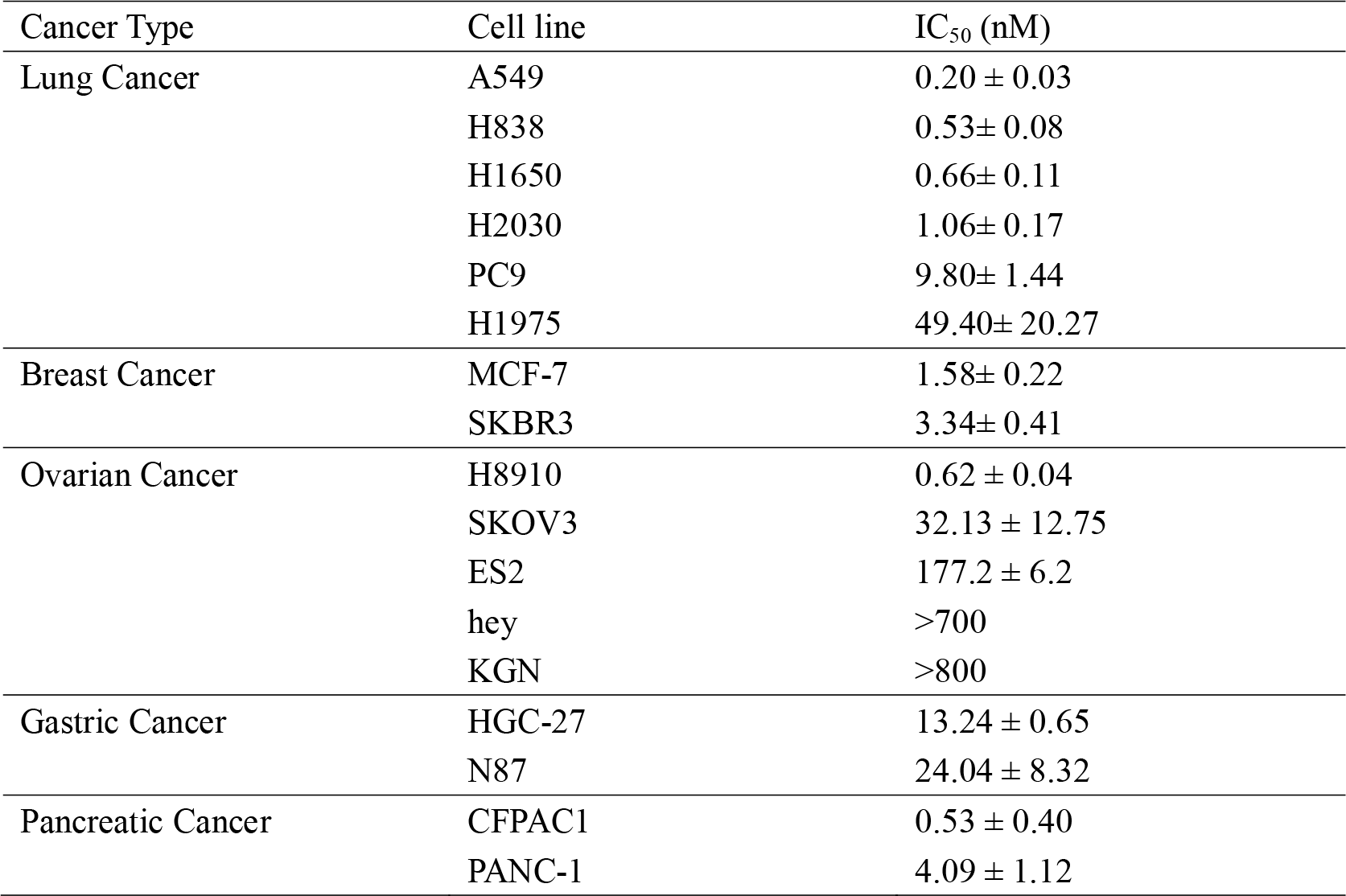
IC_50_ values of 16A-MMAE for cancer cell lines.

### In vivo antitumoral activity of 16A-MMAE

The *in vivo* antitumoral activity of 16A-MMAE was evaluated in a H838 mouse xenograft model. The results clearly showed that 16A-MMAE inhibited tumor growth in mice, in a dose-dependent manner. Tumor could be cured by two doses of 16A-MMAE at 3 mg/kg (Fig. 6A). The minimal effective dose was 1 mg/kg by two doses (Fig. 6B) or single dose (Fig. 6C).

**Fig 6.**
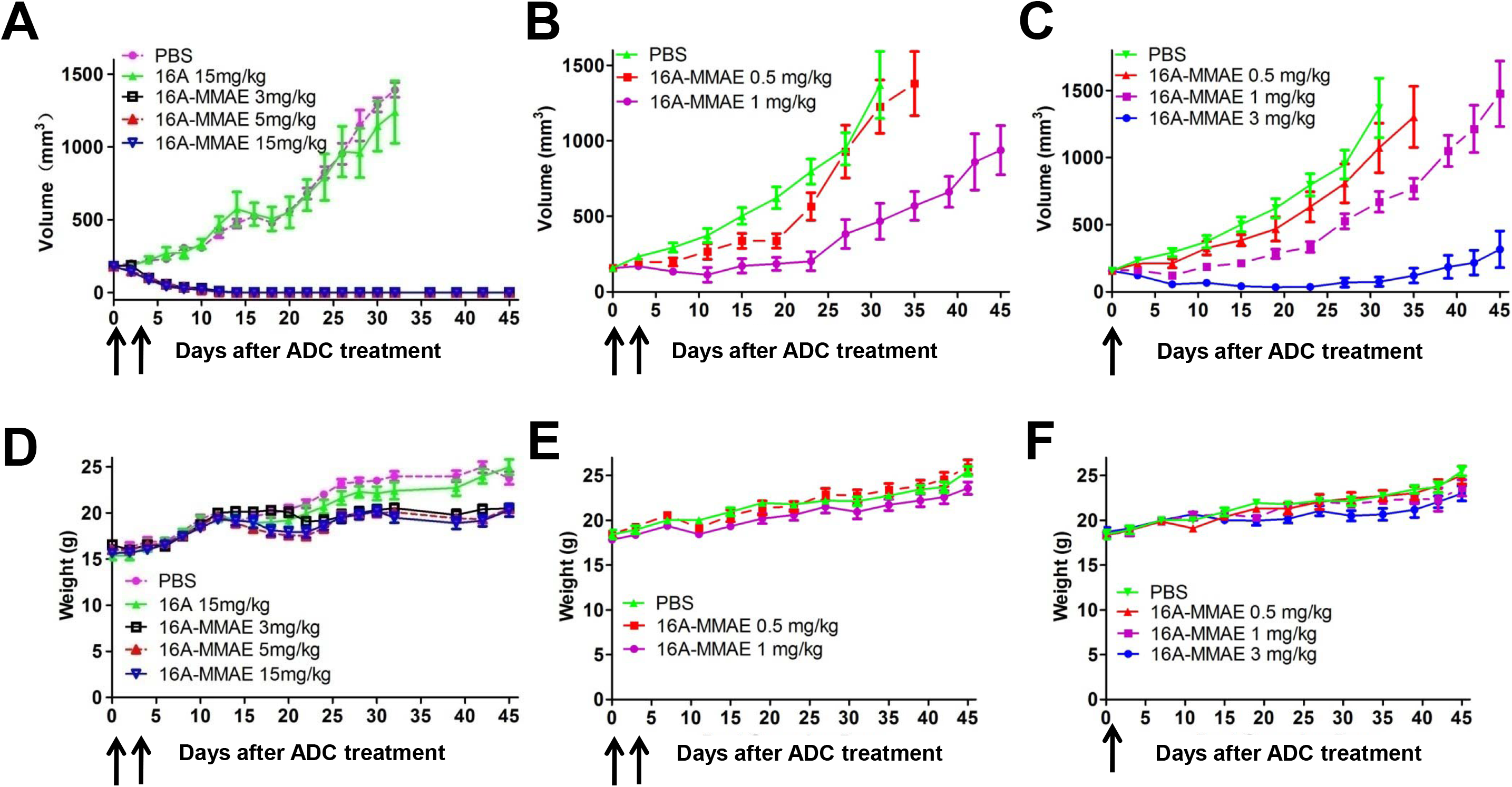
Antitumoral effect of 16A-MMAE in H838 xenograft model. **(A)** Groups of mice (Balb/c nu/nu, female, 6-week-old) bearing ~200 mm^3^ H838 tumor were treated with 16A-MMAE or non-conjugated antibody at 3 mg/kg, 5 mg/kg or 15 mg/kg, respectively. Arrows indicate time of drug treatment. **(B,C)** Mice bearing ~200 mm^3^ H838 tumor were treated with 16A-MMAE (one or two doses) at 0.5 mg/kg, 1 mg/kg or 3 mg/kg, respectively. Arrows indicate time of drug treatment. **(D,E,F)** The body weight of treated mice was measured in all groups. Data points represent mean ± SEM (n = 5).

### In vivo toxicity of 16A-MMAE

The *in vivo* toxicities of the 16A-MMAE in tumor-bearing mice were monitored by body weight. No significant changes were observed in treated groups under the dose of 5 mg/kg as compared to PBS control (Fig. 6D, 6E and 6F). To better assess drug toxicities, hMUC1 transgenic mice^22^ (024631-C57BL/6-Tg(MUC1)79.24Gend/J, from Jackson Laboratory) were administered by escalating doses of 16A-MMAE (0, 3, 15 and 30 mg/kg) and sacrificed at 3, 14 or 28 days after drug treatment. Tissue sections of the major organs (heart, liver, spleen, lung, kidney, stomach, intestine and pancreas) were analyzed after H&E staining. No obvious pathologic changes were found at above three time points for the group treated at the dose of 3 mg/kg. For the 15 and 30 mg/kg groups, minor pathologic changes were observed in some tissues (liver, lung and kidney, Fig. S7B and 7C). Immunohistochemical staining of normal hMUC1-Tg mice tissues by 16A antibody showed that the target organs of toxicity are partly associated with the antibody binding. The 16A antibody binding was observed not only in liver, lung, and kidney, but also in stomach, colon, cecum, rectum, salivary gland, trachea, uterus, and testis. No staining by 16A antibody was found in the heart, spleen, duodenum, jejunum, ileum, esophagus, brain, adrenal gland, sternum, vagina, oviduct, ovary, skin, bladder, bicipital muscle, epididymis, prostate and seminal vesicle (Fig. S8).

The *in vivo* antitumoral activity of 16A-MMAE was also tested by a syngeneic tumor model in hMUC1-Tg mice using a B16-OVA cell line stably transfected by human MUC1 gene. The results showed that 16A-MMAE inhibited tumor growth in mice, in a dose-dependent manner (Fig. S9). Tumor could be inhibited by two doses of 16A-MMAE at 10 mg/kg.

## Discussion

Tissue positivity is critical for developing antibody-based therapeutics. The potential of post-translational modification of glycoproteins has gain recent recognition in field of cancer therapy. For example, the most common glycan structures in lung cancer include Tn antigen (GalNAc), STn antigen (Neu5Acα2-6GalNAc), and ST antigen (NeuAcα1-3Gaβ1-3GalNAc).^28^ However, the exact glycopeptide sequences in every individual cancer patients are highly variable and diverse, due to the “assembly line” nature of glycosylation pathway.^29^ Big data on the glycopeptide epitopes caused by such abnormal glycosylation on specific glycoproteins in cancer population is unavailable. We previously predicted the glycopeptide sequences in lung cancer by MATLAB software.^21^ However, these sequences have not been verified by mass spectrometry analysis. The availability of monoclonal antibodies specific to synthetic glycopeptides partially addresses this problem as immunohistochemical staining by such monoclonal antibodies support the expression of a glycopeptide sequence, in spite of the caveat that the respective monoclonal antibody may have cross reactivity to structurally related other glycopeptide sequences.

Our results showed that 16A mAb, which targets the GSTA motif of MUC1,^21^ broadly binds to multiple types of cancer, including the triple negative breast cancer and gastric cancer. The positivity of 16A mAb staining in breast cancer (90%) is higher than SAR566658,^30, 31^ which binds to a sialylated unknown MUC1 sequence in bladder, breast, ovary, pancreatic, head and neck cancers with positivity of 59%, 29-35%, 70%, 59%, and 17% respectively. For those cancer tissues negative by 16A mAb staining, other mAbs targeting PDTR or GVTS motifs are worthy of further investigation.

Several ADCs targeting MUC1 have been reported. Lovat group reported the efficacy of hHuHFMG1 antibody-drug conjugate in esophageal adenocarcinoma.^32^ Kufe group reported an ADC targeting non-glycosylated C-termial of MUC1.^33^ The antitumoral efficacy and the toxicity are two major criteria to be studied for their clinical application. A phase I clinical trial for SAR566658 in patients with CA6-positive ovarian, pancreatic, and breast tumors has been completed.^30, 31^ SAR566658 is CA6 mAb conjugated to DM4, a maytansinoid derivative, by SPDB (N-succinimidyl-4-(2-pyridyldithio) butanoate) linker, a hindered disulfide bond stable linker which is stable in blood stream. Partial response was observed in breast, ovarian and lung cancers. Dose-limiting toxicities were observed at 240 mg/m^2^ (diarrhea and keratitis). Late occurrence of reversible corneal toxicity was observed at > 150 mg/m^2^.

In this study, we used hMUC1 transgenic mice model for further assessing the toxicity of 16A-MMAE. Our safety data showed that 16A-MMAE was well-tolerated in hMUC1 transgenic mice at 3 mg/kg. At higher dose, 16A-MMAE exhibited a dose-limiting toxicity. However, the tissue toxicity is not clearly associated with 16A mAb positivity. The mechanism of toxicity remain to be studied. It might be due to the low level of expression of 16A epitopes in target organs. Alternatively, the toxicity might be due to the non-specific release of MMAE to mouse serum, as we used ValCit linkers which can be hydrolyzed in mouse plasma by carboxylesterase 1c.^24,34^

Drug pay load is another critical factor for the toxicity of ADC. Recently, topoisomerase I inhibitor payload showed impressive success in clinical trials, including trastuzumab deruxtecan (Her2/Exatecan Antibody-Drug Conjugate),^35^ Labetuzumab Govitecan (CEACAM5/SN-38 Antibody-Drug Conjugate),^36^ Sacituzumab Govitecan (Trop-2/SN-38 Antibody Drug Conjugate).^37^ Such moderately cytotoxic pay loads often led to no-observed-adverse-effect level (NOAEL) in dose escalating studies, thus serve as promising conjugates to be tested for mAbs specific to cancer glycopeptides. Novel conjugation methods which are more specific and potent in conjugating drug payloads to antibodies are also evolving.^38^ More interestingly, non-internalising ADC that releases its drugs upon a click reaction with a chemical activator was recently developed, which showed efficacy in treating cancer that are resistant to those ADCs must be internalized.^39^

In summary, this study reports a neo-antigen epitope generated by abnromal post-translational modification of glycoproteins, with high positivity in a broad spectrum of cancer types. The balance of antitumoral efficacy and toxicity can further be fine-tuned by optimizing the linker and the drug payload.

## Author contributions

Dapeng Zhou, Patrick Hwu and Wei Huang designed this study. Deng Pan, Yubo Tang, Jiao Tong, ChengmeiXie, Jiaxi Chen, Chunchao Feng, Patrick Hwu and Wei Huang contributed to the collection, analysis and interpretation of data. Deng Pan and Dapeng Zhou wrote the manuscript. All authors read and approved the final manuscript.

