## Supplemental Online Materials for "An antibody drug conjugate targeting a GSTA glycosite-signature epitope of mucin1 expressed by non-small cell lung cancer": Supporting_material.docx

**Supplemental Online Figure Legends:**

**Figure S1: IHC staining of tissues from lung cancer patients by 16A antibody.**

Lung cancer tissue slides were from Pantomics (Richmond, CA). Slides were stained as described in the text by Crownbio, China.

**Figure S2: IHC staining of tissues from breast cancer patients by 16A antibody.**

Breast cancer tissue slides were from Pantomics (Richmond, CA). Slides were stained by Crownbio, China.

**Figure S3: IHC staining of gastric, colon and rectum cancers by 16A antibody.**

Cancer tissue slides were from Pantomics (Richmond, CA). Slides were stained by Crownbio, China.

**Figure S4: IHC staining of paired lung cancer and peritumoral tissues by 16A antibody.**

Slides containing paired lung cancer and peritumoral tissues were from Pantomics (Richmond, CA). Slides were stained by Crownbio, China.

**A),B)** Human tissue array containing multiple tissues from the lung adenocarcinoma and squamous carcinoma patients were stained by 16A antibody. a1 and a2 (b1 and b2) are tumor tissue, a3 (b3) is the same patient’s peritumoral tissue.

**Figure S5: IHC staining of tissues from healthy individuals by 16A antibody.**

Slides containing multiple types of tissues from healthy individuals were from Pantomics (Richmond, CA). Slides were stained by Crownbio, China.

**Figure S6: *In vitro* antitumoral efficacy of 16A-MMAE**

Up panels: inhibition of cancer cell lines by increasing concentrations of 16A-MMAE.

Lower panels: flow cytometry staining of cell lines by 16A antibody.

**Figure S7: Toxicity of 16A-MMAE in hMUC1 transgenic mice.**

16A-MMAE was administered to hMUC1 transgenic mice (n = 6 per group, three males and three females) via tail vein injection at a single dose of 0, 3, 15, or 30 mg/kg. Tissues were harvested for clinical pathological assessment on days 3, 14, and 28 (two mice per group at each time point, one male and one female). Histopathological changes of heart, liver, spleen, lung, kidney, gastric, pancreatic, and small intestine were examined after H&E staining (original magnification ×100).

**Figure S8: IHC staining of multiple organs of the hMUC1 transgenic mice and wild type control by 16A antibody.** Transgenic (hMUC1) and wild type mice tissues (heart, liver, spleen, lung, kidney, pancreas, stomach, duodenum, jejunum, ileum, colon, cecum, rectum, esophagus, brain, salivary gland, trachea, adrenal gland, sternum, vagina, oviduct, ovary, uterus, skin, eyeball, bladder, bicipital muscle, epididymis, testis, prostate, and seminal vesicle) were stained by 16A antibody (original magnification ×100).

**Figure S9. Antitumoral effect of 16A-MMAE in B16-OVA-hMUC1 model.** The *in vivo* antitumoral activity of 16A-MMAE was evaluated in a B16-OVA-hMUC1 cell transplant model *in vivo* using C57BL/6-Tg(MUC1)79.24Gend/J mice. 16A-MMAE inhibited tumor growth in mice in a dose-dependent manner. Tumor could be inhibited by two doses of 16A-MMAE at 10 mg/kg.

Figure S1

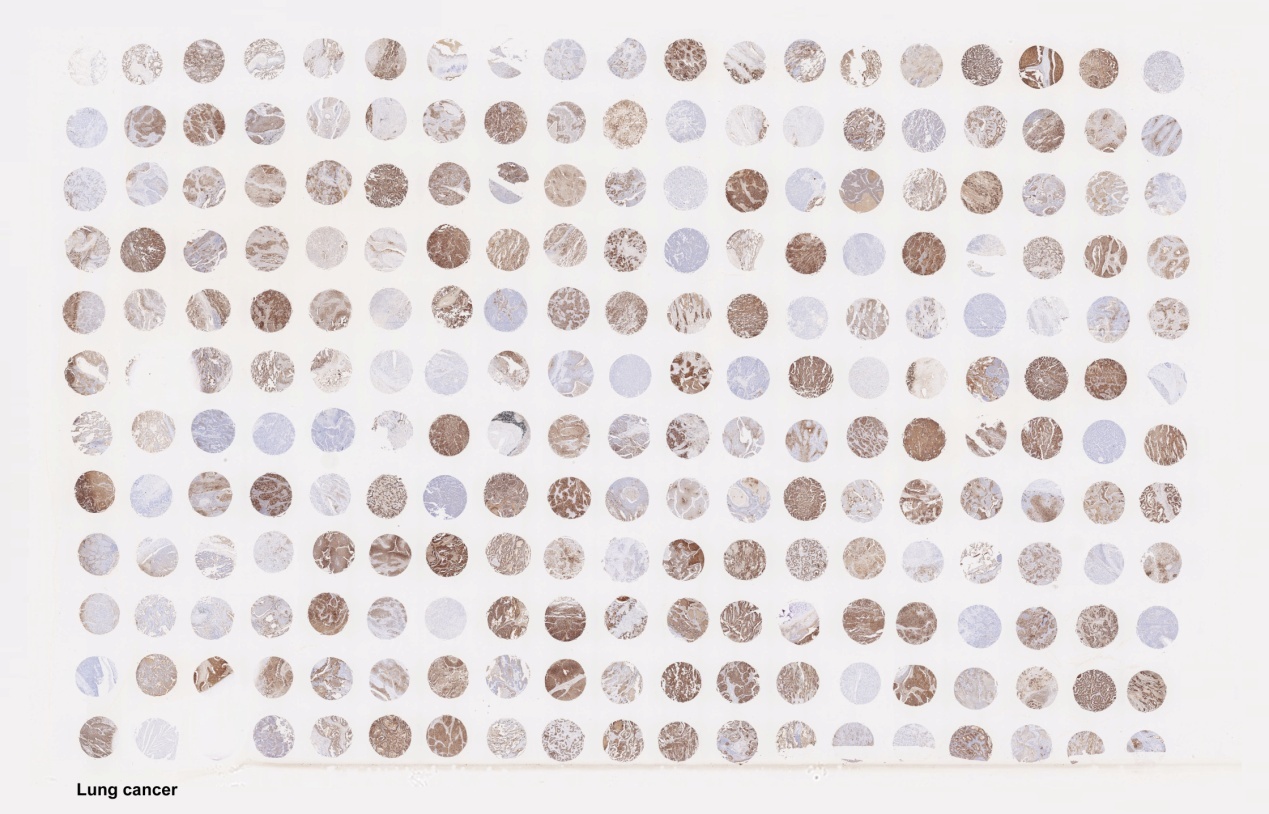

Figure S2

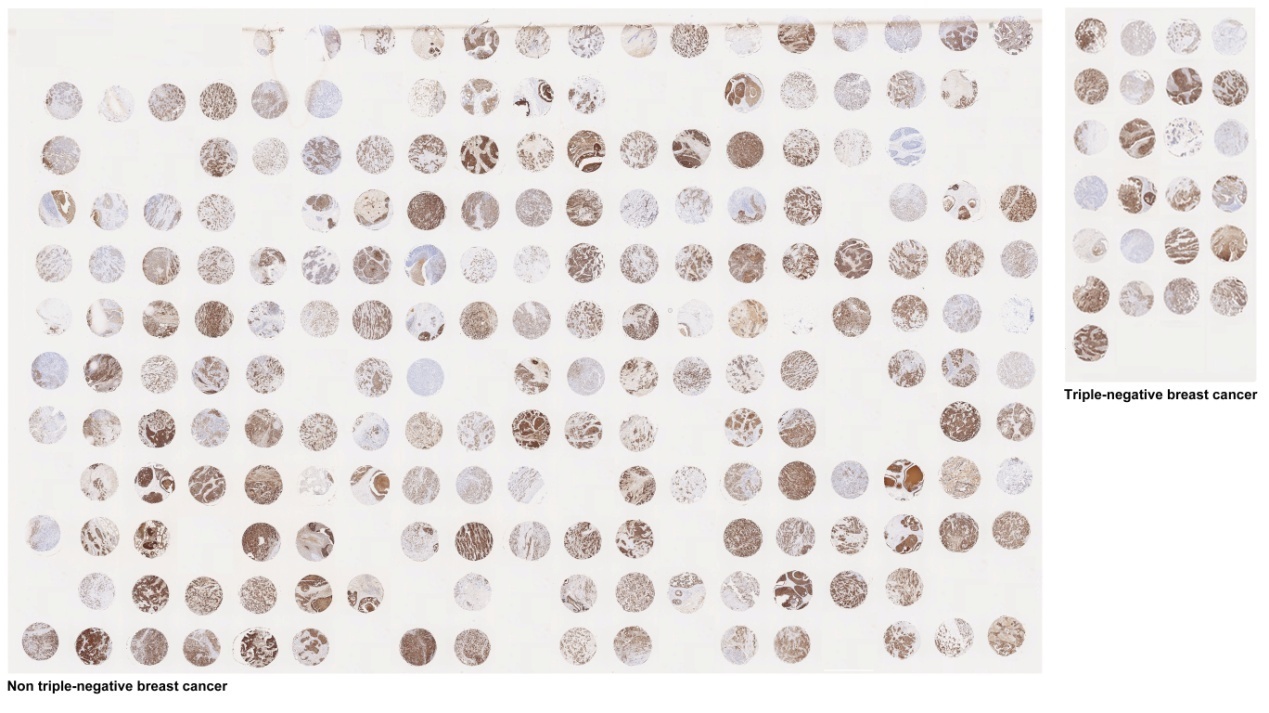

Figure S3

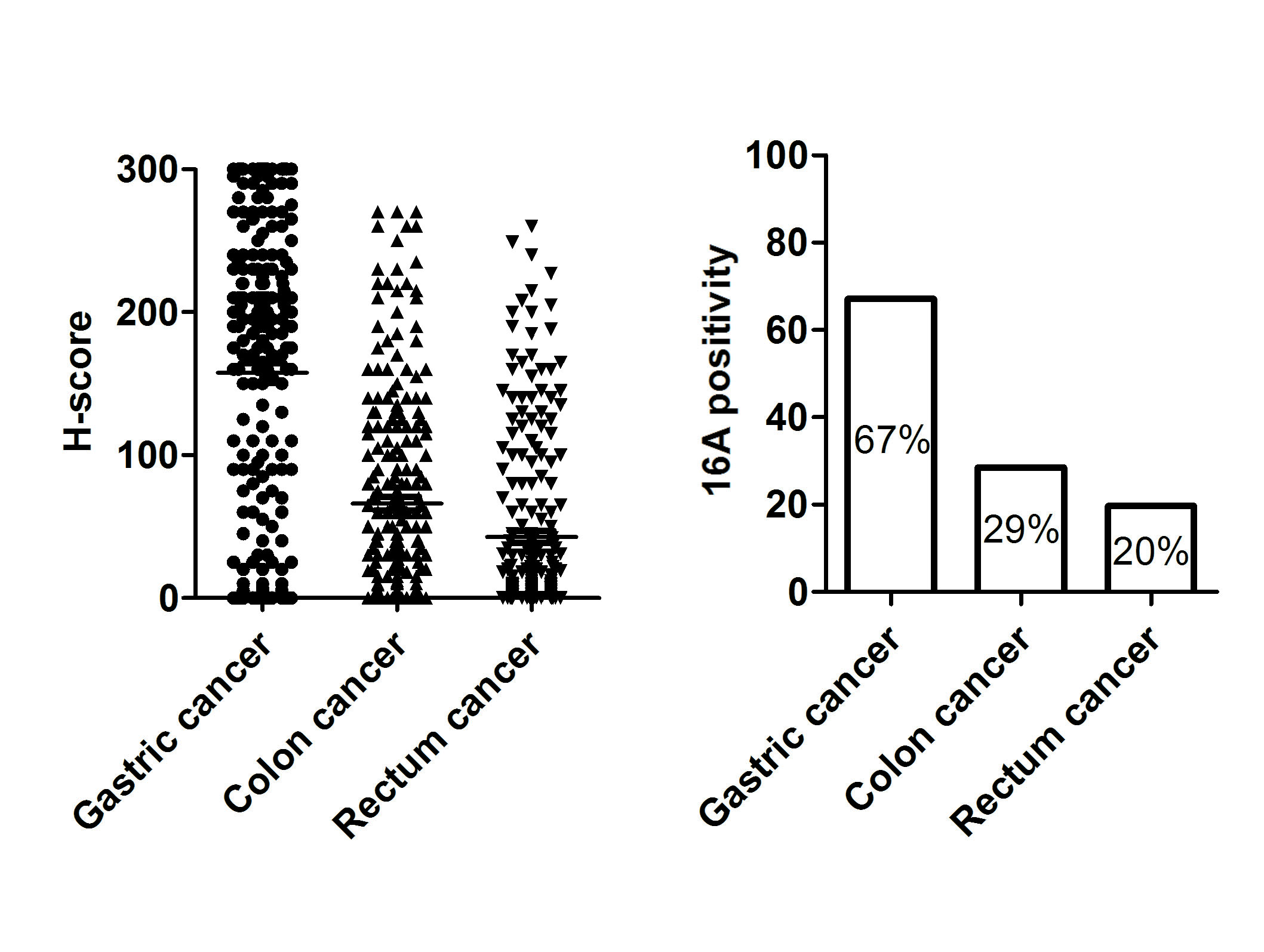

Figure S4

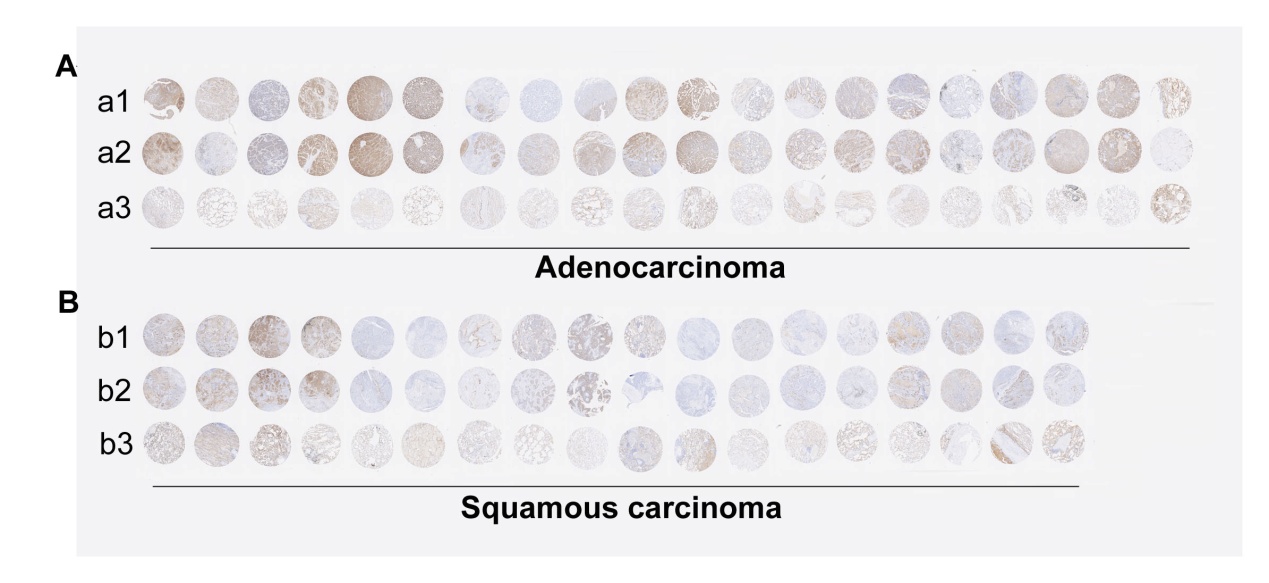

Figure S5

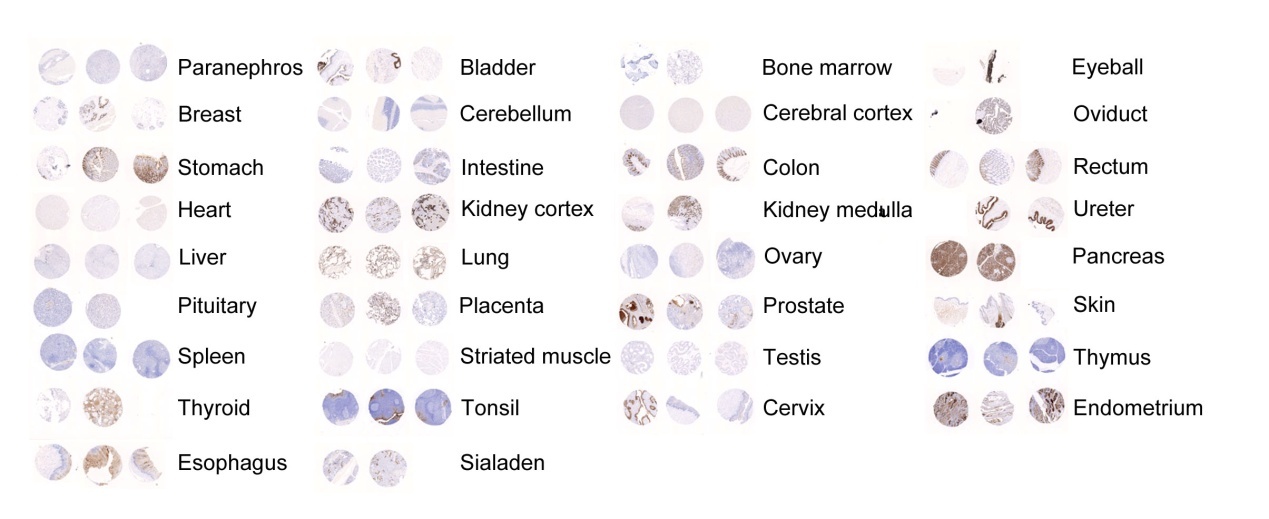

Figure S6

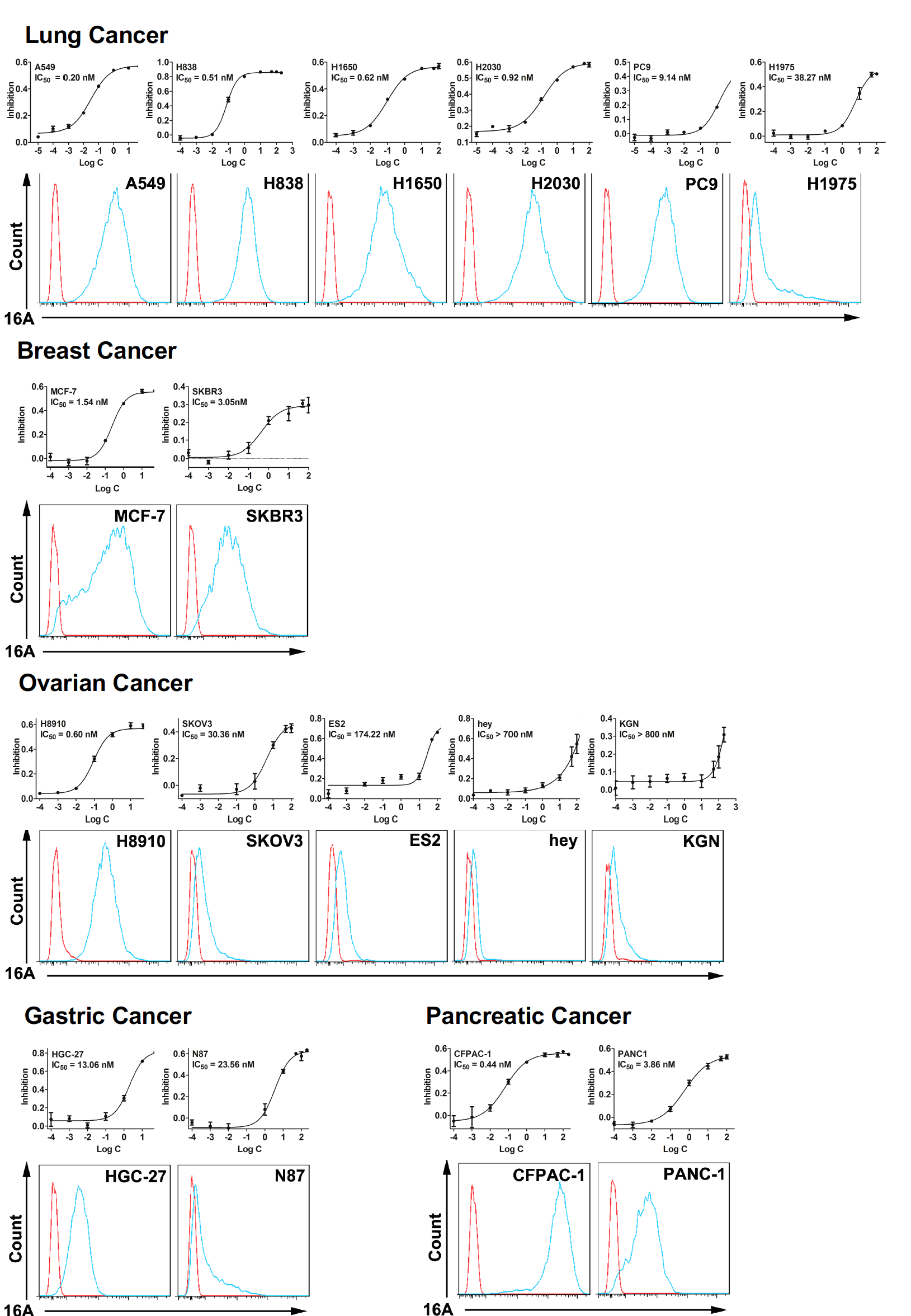

Figure S7A

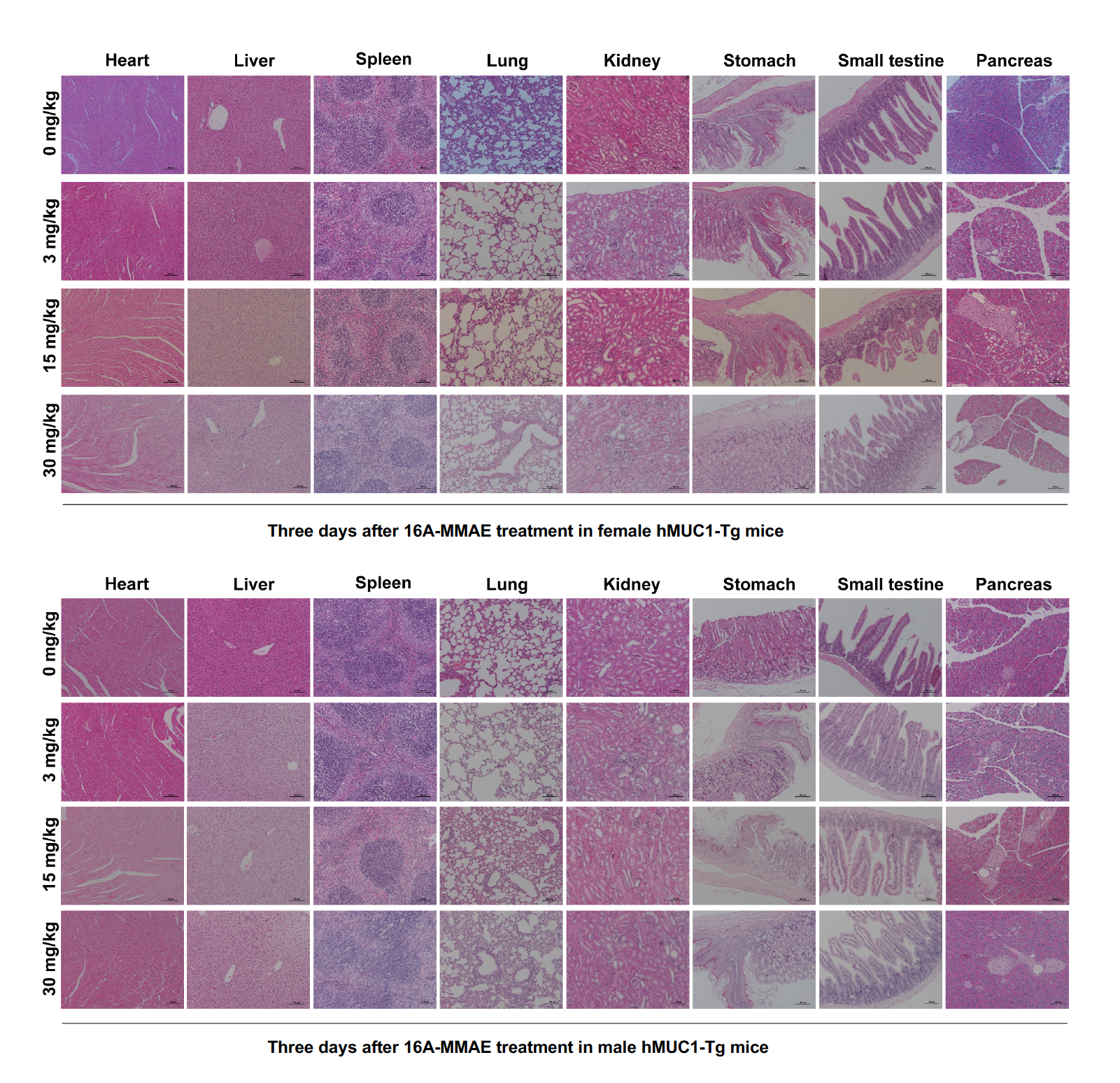

Figure S7B

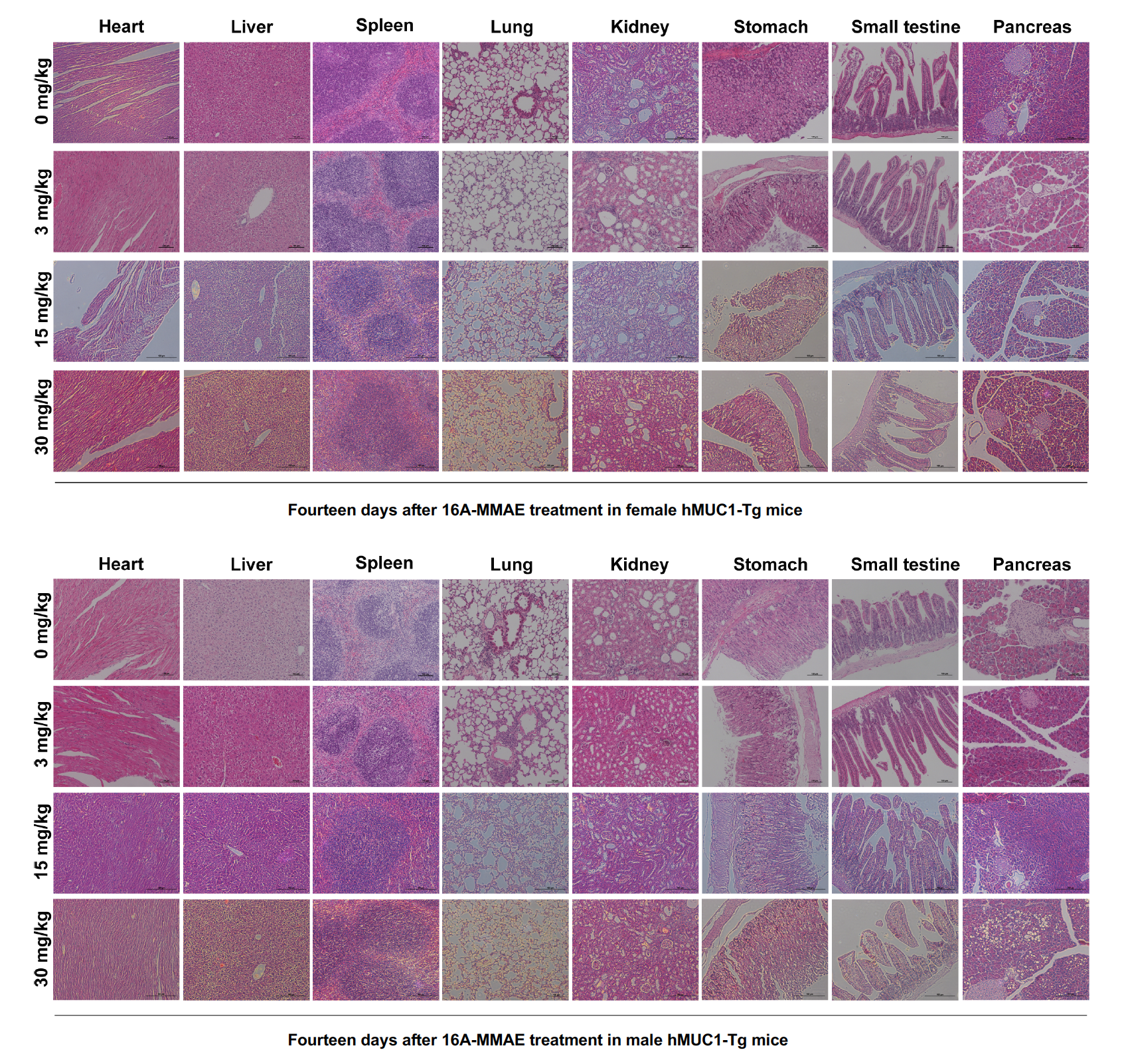

Figure S7C

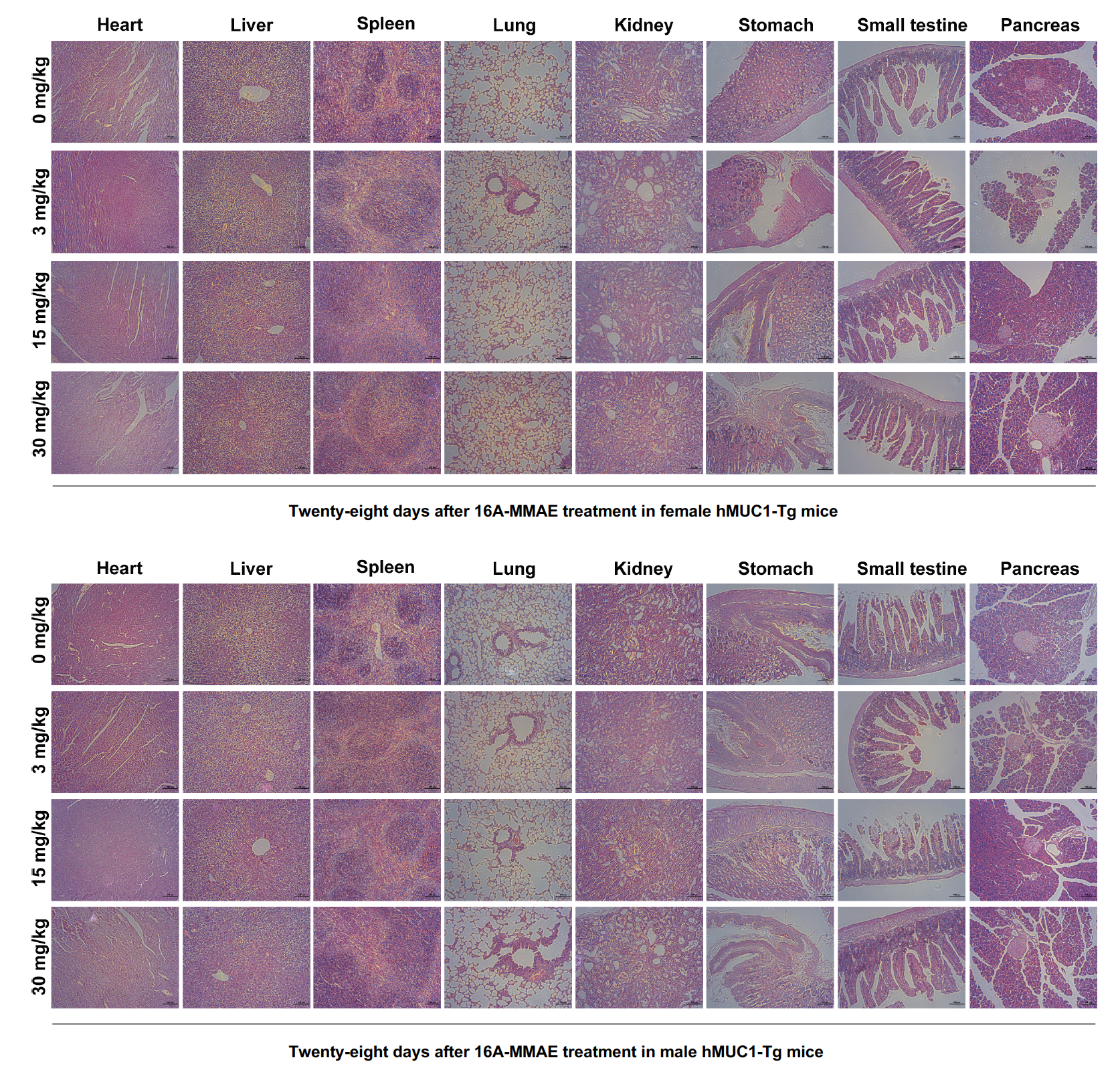

Figure S8

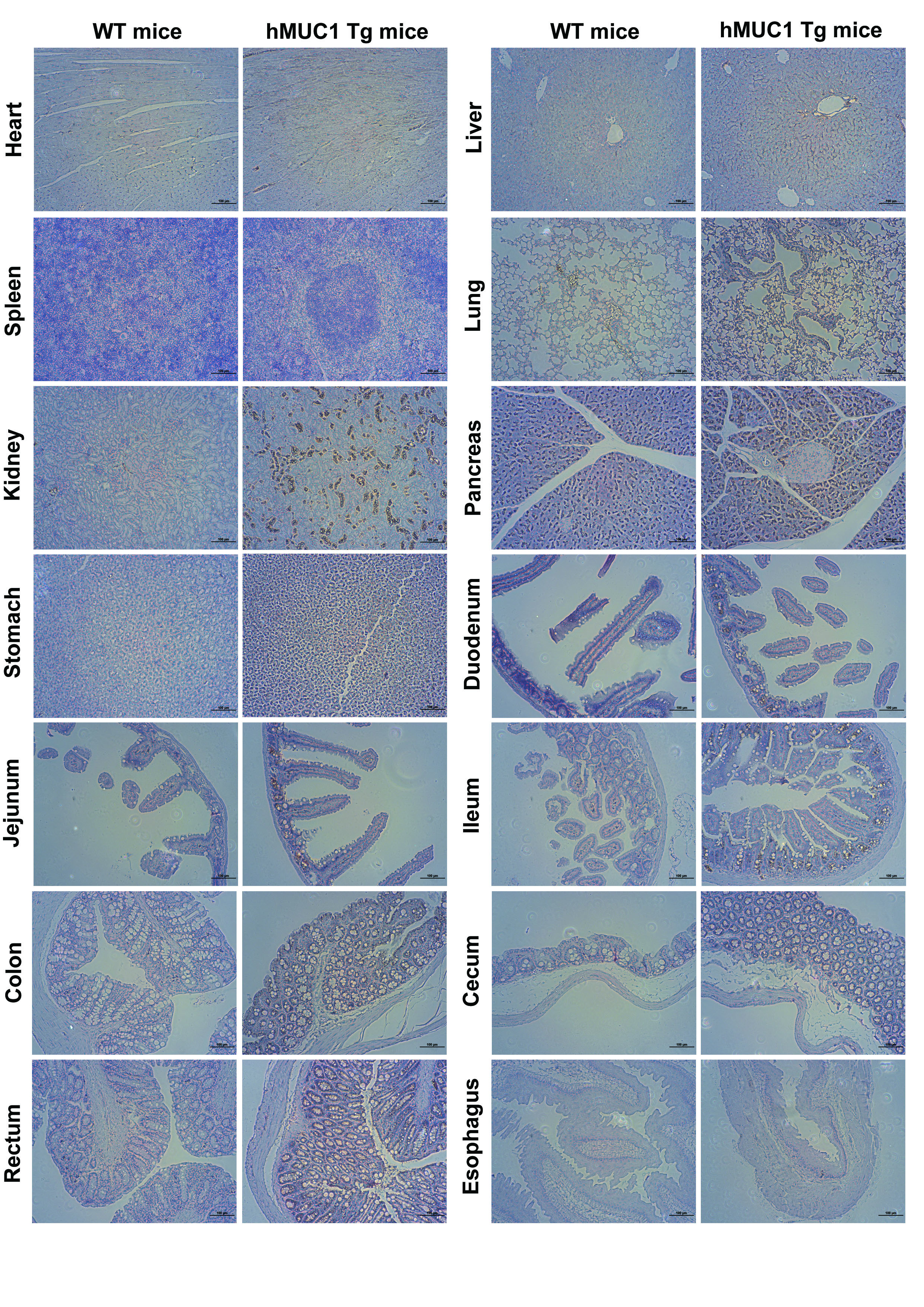

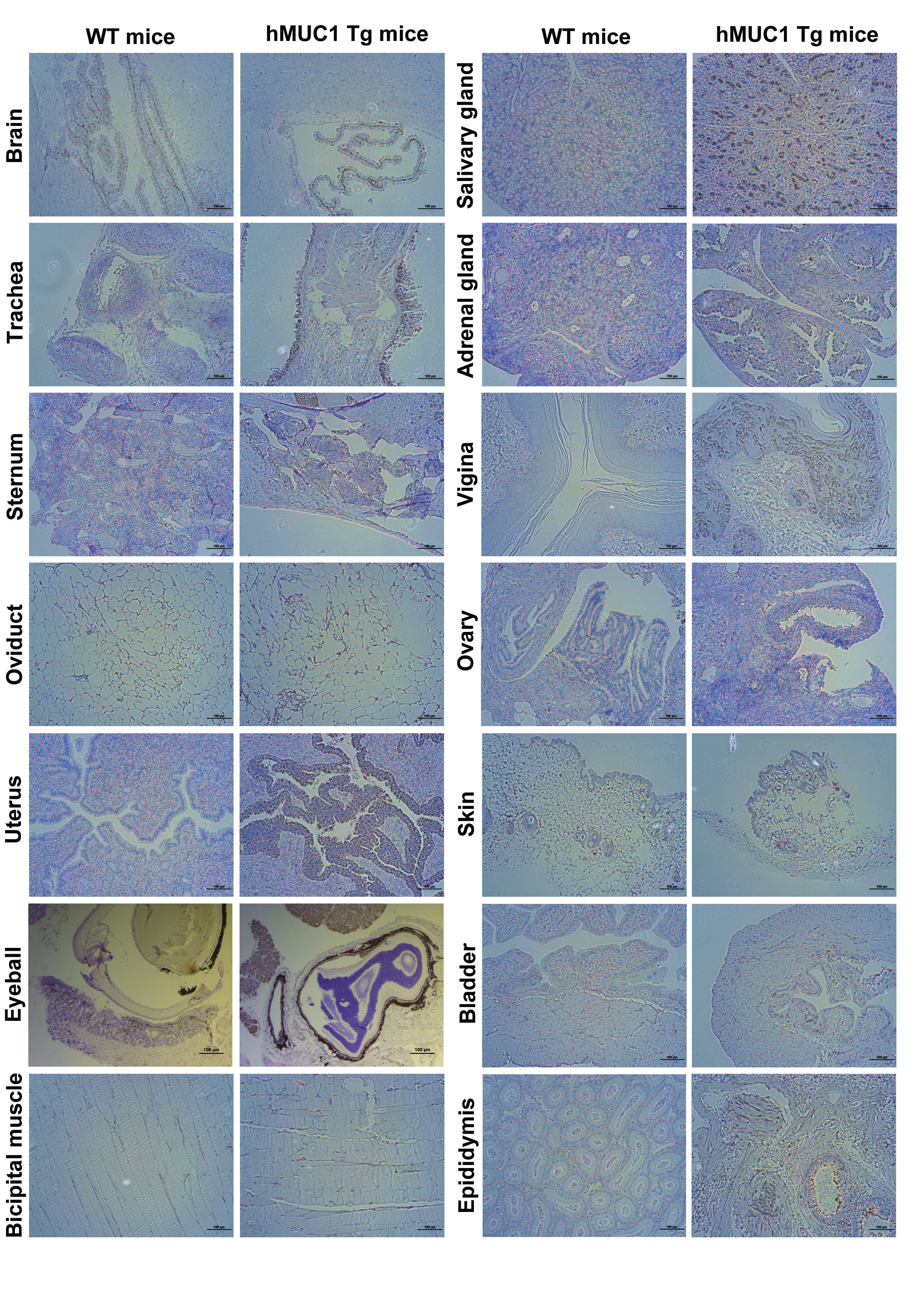

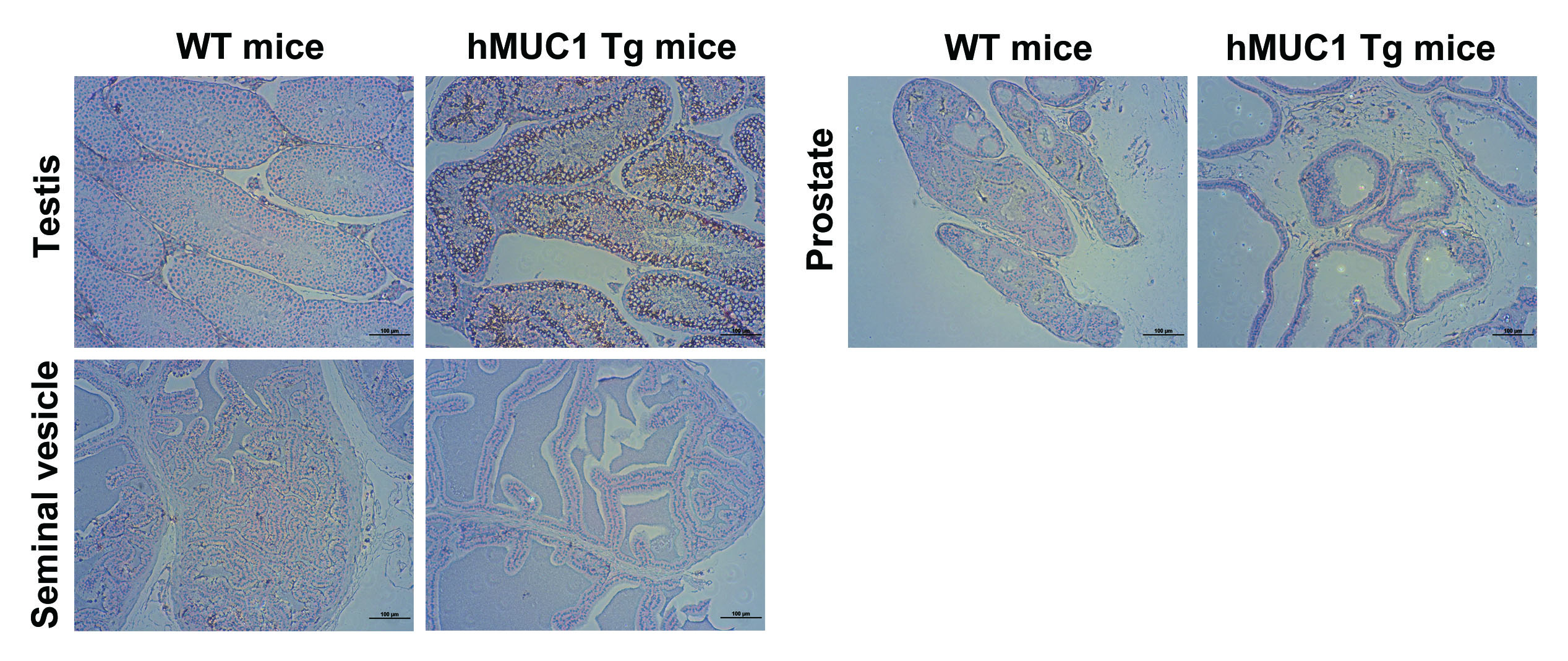

Figure S9

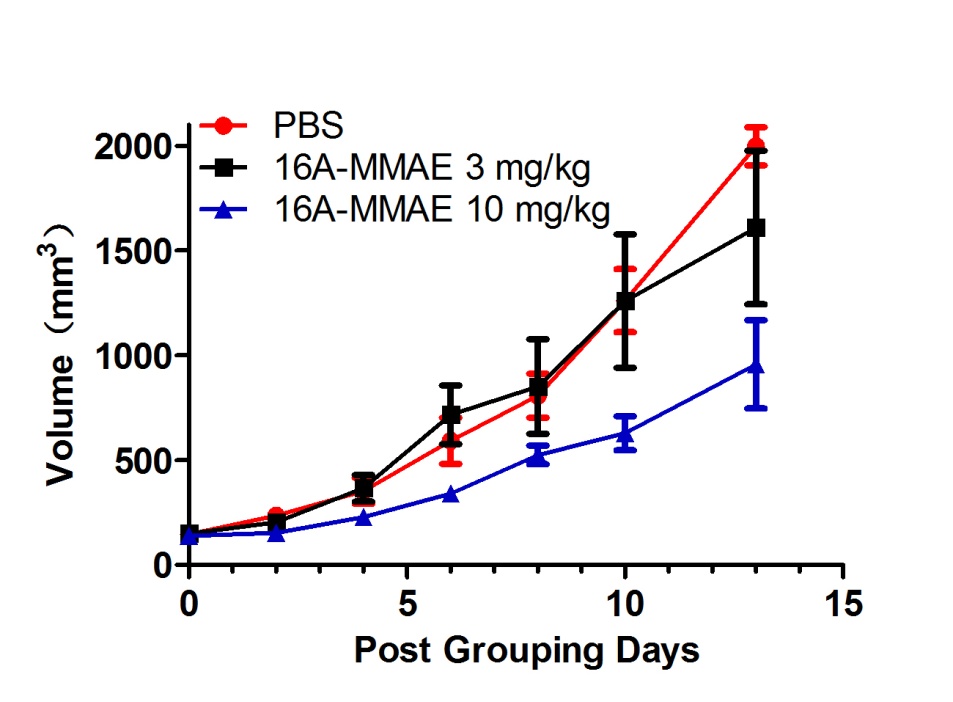

**Table S1. Pharmacokinetics of 16A-MMAE antibody-drug conjugate.**

| Parameters | 16A | 16A-MMAE |
| --- | --- | --- |
| t_1/2_ (h) | 207.00 | 144.22 |
| CL (ml/h/kg) | 0.27 | 0.36 |
| MRT (h) | 278.87 | 198.98 |

t_1/2_: Elinination Half-life.

CL: Clearance.

MRT: Mean Residence time.
